# Ultralow background membrane editors for spatiotemporal control of lipid metabolism and signaling

**DOI:** 10.1101/2023.08.31.555787

**Authors:** Xiang-Ling Li, Reika Tei, Masaaki Uematsu, Jeremy M. Baskin

## Abstract

Phosphatidic acid (PA) is a multifunctional lipid with important metabolic and signaling functions, and efforts to dissect its pleiotropy demand strategies for perturbing its levels with spatiotemporal precision. Previous membrane editing approaches for generating local PA pools used light-mediated induced proximity to recruit a PA-synthesizing enzyme, phospholipase D (PLD), from the cytosol to the target organelle membrane. Whereas these optogenetic PLDs exhibited high activity, their residual activity in the dark led to undesired chronic lipid production. Here, we report ultralow background membrane editors for PA wherein light directly controls PLD catalytic activity, as opposed to localization and access to substrates, exploiting a LOV domain-based conformational photoswitch inserted into the PLD sequence and enabling their stable and non-perturbative targeting to multiple organelle membranes. By coupling organelle-targeted LOVPLD activation to lipidomics analysis, we discovered different rates of metabolism for PA and its downstream products depending on the subcellular location of PA production. We also elucidated signaling roles for PA pools on different membranes in conferring local activation of AMP-activated protein kinase signaling. This work illustrates how membrane editors featuring acute, optogenetic conformational switches can provide new insights into organelle-selective lipid metabolic and signaling pathways.

**TOC Graphic:** 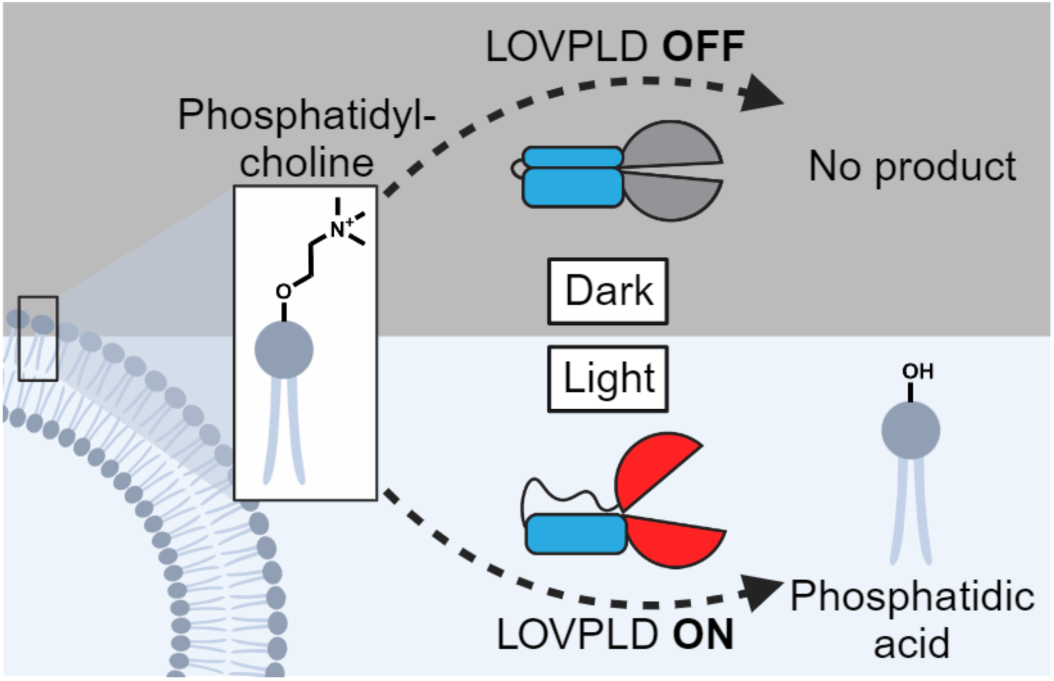

## Introduction

Lipids are remarkably pleiotropic biomolecules, mediating myriad cellular signaling events beyond their key structural roles within organelle membranes^1–3^. However, the multifunctionality of lipids also makes individual lipid signaling pathways difficult to unravel. A prime example of a pleiotropic lipid is phosphatidic acid (PA). In mammalian cells, PA biosynthesis occurs via at least three main pathways: acylation of lysophosphatidic acid (LPA) by lysophosphatidic acid acyltransferases (LPAATs), phosphorylation of diacylglycerol (DAG) by DAG kinases (DGKs), and hydrolysis of phosphatidylcholine (PC) by phospholipase Ds (PLDs)^4–6^. PA in turn serves as a key precursor for the biosynthesis of most other phospholipids and triglycerides^7^.

PA metabolism is highly regulated and dependent upon both inter-organelle lipid transport mediated by lipid-transfer proteins and lipid-metabolizing enzymes^8^. For example, the lipid transfer protein Nir2 facilitates PA transport from the plasma membrane to the endoplasmic reticulum (ER), where it is converted to CDP-DAG, a key intermediate in phosphatidylinositol and phosphatidylglycerol biosynthesis^9^. PA also participates in several major signaling pathways, either indirectly by conversion to other signaling molecules such as LPA^10^, or by directly binding to effector proteins, such as Large tumor suppressor kinase 1 (LATS) in Hippo signaling and Liver kinase B1 (LKB1) in AMP-activated protein kinase (AMPK) signaling^8,11–13^. Additionally, PA can function as a membrane anchor for recruiting protein complexes to specific organelle membranes, indicating that the subcellular localization where pools of PA are produced have a functional importance^14–17^.

To elucidate physiological functions of PA pools on different organelles, approaches for rapid addition or depletion of PA from cellular membranes are needed. Genetic perturbations such as knockout, knockdown, or overexpression of lipid-modifying enzymes are problematic because they occur on long, multi-day timescales. Whereas pharmacological inhibitors and bulk lipid dosing to cells act rapidly, they typically lack spatial control, thus making them not ideally suited for studying localized effects of lipid perturbations^3,8,18^. To address this technological gap, we have previously developed an optogenetic strategy for rapid, blue light-controlled production of PA on target organelle membranes in mammalian cells^19^. Our optogenetic PLD (optoPLD), harnesses the CRY2–CIBN heterodimerization system to enable rapid recruitment of a bacterial PLD (from *Streptomyces* sp. strain PMF) that is orthogonal to the human proteome to produce PA from PC, the most abundant phospholipid in cell membranes, on target membranes of interest. In addition to catalyzing PC hydrolysis to produce PA, PLDs also mediate transphosphatidylation with primary alcohols to produce a variety of phosphatidyl alcohols, making these enzymes a useful starting point for a general phospholipid editor.

Most recently, we substantially improved the catalytic activity of optoPLD to yield a family of super-active PLDs (superPLDs) by increasing its stability in the reductive cytosolic environment via mammalian cell-based directed evolution^20^. SuperPLDs catalyze both hydrolysis and transphosphatidylation with significantly higher efficiency than wild-type PLD^PMF^, indicating their potential as highly versatile phospholipid editors within living cell membranes. However, optogenetic superPLDs exhibited increased background activity in the absence of the light trigger, i.e., without membrane recruitment, limiting their temporal resolution and causing unwanted, off-target effects. This inherent problem likely arises from stochastic encounters of cytosolic superPLDs with membranes via diffusion in the dark. Most critically, this background PA production led to some cytotoxicity in mammalian cells transiently transfected with certain optogenetic versions of the most potent superPLDs even when kept in the dark. Signaling and biosynthetic pathways can also become altered in cells expressing optogenetic superPLDs as they acclimate to chronically high levels of PA, thus making them less viable as tools to probe PA signaling under physiological conditions.

Herein, we present an ultralow background membrane editor capable of producing PA with high spatiotemporal control that uses a fundamentally different approach for turning PA production on and off. Instead of controlling the recruitment of an always-on superPLD to the substrate, i.e., to organelle membranes, we devised a strategy for directly controlling the enzymatic activity of superPLD. Toward this goal, we exploited the light-oxygen-voltage (LOV) 2 domain of phototropin 1 derived from *Avena sativa* (AsLOV2), which changes conformation upon blue light illumination. With the expectation that the light-induced conformational change of the AsLOV2 domain could impact the superPLD structure and consequently its activity, we grafted an engineered variant of AsLOV2 domain^21^ into a flexible loop of superPLD. The resulting photoswitchable PLD, which we have termed LOVPLD, displayed ultralow background activity, no discernable cytotoxicity, and compatibility with multiple organelles and cell lines. We demonstrate the utility of LOVPLD by addressing two important questions in PA metabolism and signaling: first, to elucidate differential turnover rates of PA on distinct organelle membranes, and second, to investigate acute effects of PA on AMPK signaling. As such, LOVPLD represents a powerful addition to the toolbox for the study of PA signaling, and we envision that our strategy to control enzymatic activity, not merely localization, of heterologous enzymes can be broadly applicable to create additional biomolecular editors for studying not only lipid biology but also other metabolic signaling pathways.

## Results

### LOV domain insertion into superPLD renders it photoactivatable

To directly control superPLD activity, we sought to insert into it a reversible conformational switch so that the activity of superPLD could be allosterically controlled by this conformational change. Whereas many conformation-switching domains exist, we chose hLOV1^21^, an engineered version of the AsLOV2 domain as the most promising candidate due to its small size, rapid activation and deactivation kinetics, tighter dark state binding, and well-documented use in optogenetic tools^22–24^. We termed the chimeric proteins resulting from the insertion as LOVPLDs (**Fig. 1A**). To maximize signal, we chose the most active superPLD (superPLD^high^, clone 2-48)^20^ as the starting point. Based on sequence conservation analysis using ConSurf^25,26^, we identified 19 positions with low evolutionary conservation located within flexible loops that we hypothesized would have a higher chance of tolerating the insertion of the hLOV1 domain **(Fig. S1)**. Finally, we designed the construct to be constitutively anchored on the plasma membrane (PM) using the first 10 amino acids of the Lyn protein (Lyn_10_)^27^ so that the enzyme would have immediate access to its lipid substrate, PC, upon photoactivation. We expressed each of these 19 PM-anchored LOVPLD variants in HEK 293T cells and analyzed their activity using a click chemistry-based, in-cell PLD activity assay termed IMPACT^28,29^ **(Fig. S2A–B)**. In IMPACT, cells are first treated with azidopropanol (AzPrOH), which is used by PLD as a transphosphatidylation substrate to produce azido phosphatidyl alcohols. The cells are then treated with a cyclooctyne-fluorophore conjugate (bicyclononyne (BCN)-BODIPY) to generate BODIPY-tagged phosphatidyl alcohols, which serve as fluorescent reporters of PLD activity, via a bioorthogonal strain-promoted azide-alkyne cycloaddition (SPAAC) reaction **(Fig. 1B)**.

**Figure 1.**
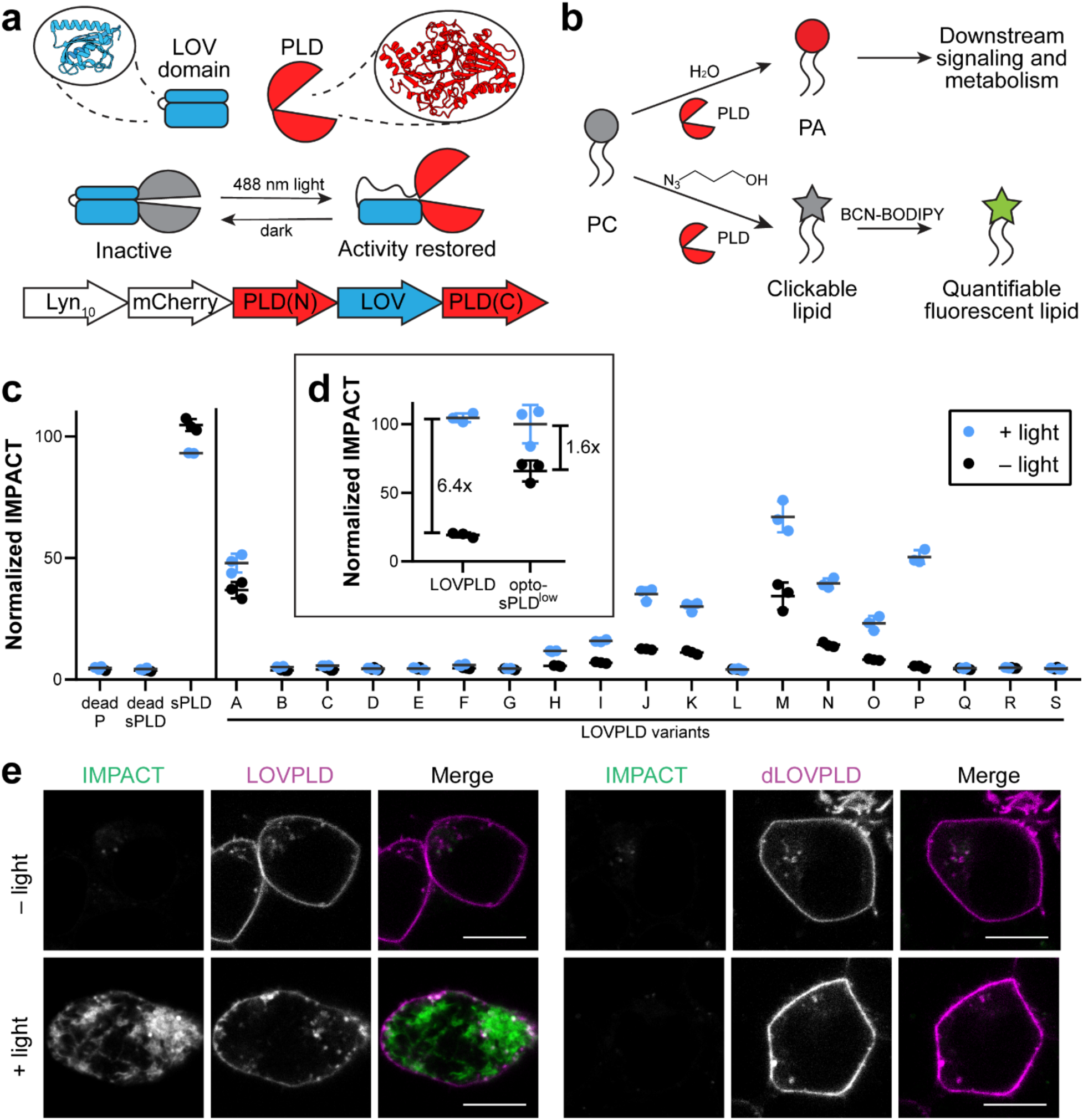
LOVPLD is a blue-light activated system for phosphatidylcholine hydrolysis and transphosphatidylation in live cells. (a) Schematic illustrating the design of LOVPLD. SuperPLD^high^ (PLD) is split into two halves (N and C) at a flexible loop, and the hLOV1 domain (LOV) is inserted at the split site. The enzyme exists in inactive conformation in the dark; upon illumination with blue (488 nm) light, the helical arm of the LOV domain relaxes, allowing the superPLD to undergo conformational changes required to gain catalytic activity. Plasmids encoding PM-anchored LOVPLD were constructed in the form of Lyn_10_-mCherry-PLD(N)-LOV-PLD(C). (b) PLD enzymes catalyze either hydrolysis of PC into PA (top) or, in the presence of primary alcohols, transphosphatidylation of PC to form phosphatidylalcohols. In the case of clickable alcohols such as 3-azidopropanol, the resultant azido phosphatidylalcohols can then be tagged with fluorescent cyclooctyne reagents (e.g., BCN-BODIPY) via SPAAC to generate fluorescent lipid reporters of PLD activity. (c) Screen of activity of 19 LOVPLD variants labeled as A to S, using IMPACT labeling, compared to controls (catalytically dead form of LOVPLD clone P (dead P), catalytically dead superPLD^high^ (dead sPLD), and constitutively active superPLD^high^ (sPLD)). IMPACT labeling was quantified as mean BODIPY fluorescence intensity measured via flow cytometry, followed by normalization to IMPACT labeling of cells expressing sPLD (n=3). (d) Comparison of IMPACT activities of LOVPLD and previously reported PM-localized optogenetic superPLD^low^ (opto-sPLD^low^) (n=3). Indicated are background-subtracted fold increase in IMPACT labeling before and after blue light illumination. (e) Representative images of IMPACT-labeled cells expressing LOVPLD and dLOVPLD. Scale bars: 10 μm.

HEK 293T cells transfected with each LOVPLD construct were exposed to intermittent blue light (5-s pulses every 1 min) for 30 min. We then measured the intensity of cellular IMPACT labeling using flow cytometry. As a control, cells transfected with a PM-localized form of the non-optogenetic parent superPLD^high^ (sPLD) were prepared, which showed similar levels of IMPACT labeling regardless of exposure to blue light. Further, catalytically dead mutants of sPLD and LOVPLD P (dead sPLD and dead P; containing the catalytic point mutation H170A) exhibited minimal IMPACT labeling, indicating that the insertion of a hLOV1 domain alone did not introduce any unexpected enzymatic activity. Of the 19 LOVPLD variants, 10 showed similar levels of IMPACT labeling as catalytically dead mutants regardless of exposure to blue light, indicating that the insertion of the hLOV1 domain at those sites destroyed enzymatic activity. However, nine variants retained activity even with the hLOV1 domain insertion, and all of these showed some light-dependent increases in IMPACT labeling **(Fig. 1C)**.

Notably, variant P showed the highest signal-to-background ratio, exhibiting near-baseline activity in the dark and ∼50% activity of the parent superPLD after illumination with blue light. We assessed the activity of LOVPLD variant P (hereafter referred to simply as LOVPLD) relative to the dimerization-based optogenetic superPLD^low^, which was previously optimized for mammalian cell-based membrane editing applications^20^. Whereas the latter achieved a 1.6-fold increase in IMPACT labeling upon blue light activation, LOVPLD reached a similar level of maximum activity but with 6.4-fold increase in IMPACT labeling, with the increase in signal-to-noise coming entirely from an almost 4-fold decrease in off-state activity **(Fig. 1D)**. We verified the PM localization of the Lyn_10_-anchored LOVPLD and catalytically dead LOVPLD (dLOVPLD) using confocal microscopy and further showed that IMPACT labeling in LOVPLD-expressing cells required blue light illumination **(Fig. 1E)**. We note that the IMPACT labeling was predominantly apparent in the ER, the major compartment to which PM-derived BODIPY-tagged phosphatidyl alcohols made by IMPACT are rapidly transported^29–32^. Collectively, these flow cytometry and imaging analyses confirm that light can effectively control the activity of LOVPLD without manipulating its localization.

A practical demonstration of the ultra-low background activity of LOVPLD is the ability to generate cell lines stably expressing the optogenetic PLD construct. Attempts using our previous dimerization-based optoPLD or optogenetic superPLDs were unsuccessful, showing inconsistent and poor expression that was lost upon successive passaging, likely due to toxicity arising from undesired PA production even when cells were kept in the dark **(Fig. S2C–D)**. In striking contrast, when we generated stable cell lines following transduction of lentiviral-encoded LOVPLD, we observed LOVPLD stable expression over multiple passages **(Fig. S2C–D)**.

### LOVPLD can be combined with induced proximity systems for finer spatiotemporal control

Although LOVPLD achieved very low background activity that would be sufficient for most biological applications, such miniscule background activity is nonetheless detectable and above baseline, i.e., extent of IMPACT fluorescence in the presence of the catalytically dead dLOVPLD. Therefore, we set out to further lower this background with a “double-gating” strategy wherein LOVPLD is combined with a blue light-induced dimerization system such that the light stimulus would serve as a single trigger for controlling both PLD catalytic activity and localization. We selected two systems: CRY2–CIBN, a popular heterodimerization pair that we used in optoPLD^33^, and improved light-induced dimer (iLID), which has much faster on-off kinetics than CRY2–CIBN^34^. We designed constructs incorporating LOVPLD into each of these systems, where one dimerization partner was fused to a PM-targeting tag (Lyn_10_ for iLID and CAAX for CRY2– CIBN), the other partner is fused to LOVPLD at either the C-terminus or N-terminus, and the two components are cloned into a bicistronic vector separated by a self-cleaving P2A peptide **(Fig. 2A)**.

**Figure 2.**
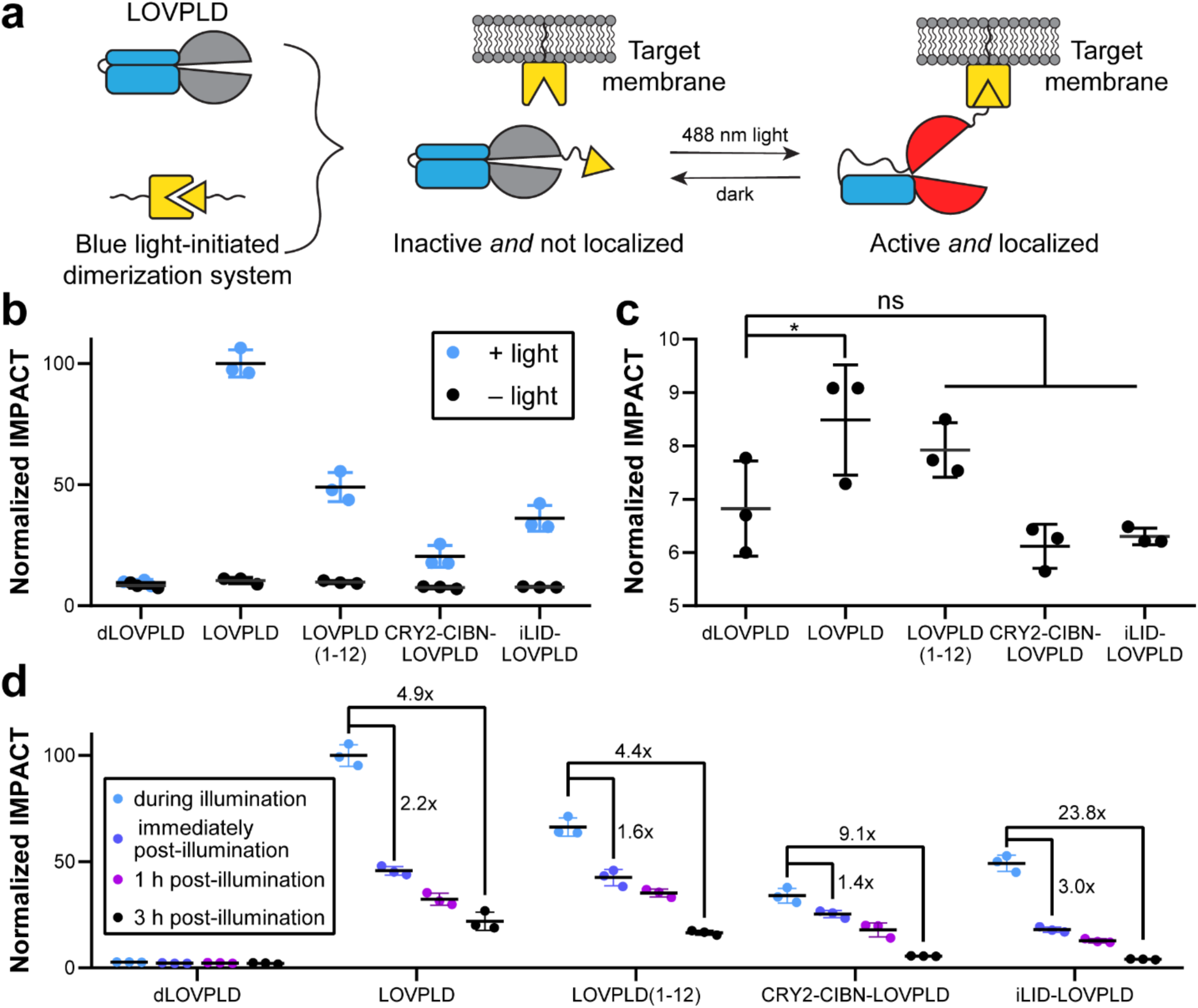
Combining LOVPLD with light-induced dimerization systems affords double-gated, ultralow background membrane editors. (a) Schematic illustrating a double-gated system featuring LOVPLD and a second blue light-initiated dimerization system. Upon blue light illumination, LOVPLD becomes activated and is translocated to the target membrane via optogenetic heterodimerization. (b) LOVPLD retains on-off activity, as assessed by IMPACT labeling, either when a different superPLD (superPLD^med^, clone 1-12) is used as the backbone (LOVPLD(1-12)) or when incorporated into a dimerization system (CRY2-CIBN-LOVPLD or iLID-LOVPLD) (n=3). (c) Zoom-in of the IMPACT labeling from (b) only in the dark, highlighting differences in background activity between the constructs. Statistical significance compared to dLOVPLD was determined by ordinary one-way analysis of variance (ANOVA) followed by Dunnett’s multiple comparisons test. *, *P* < 0.05; ns, not significant. The *P* values for the indicated comparisons to dLOVPLD are 0.0429, 0.2148, 0.5460, and 0.7574, respectively. (d) Doubly-gated LOVPLDs exhibit more rapid deactivation kinetics. IMPACT labeling of cells expressing the indicated construct either during blue light illumination, immediately after blue light illumination, or either 1 h or 3 h after returning to the dark (n=3). Indicated are background-subtracted fold decrease in IMPACT labeling immediately after blue light illumination and 3 h post-illumination.

We also designed a version of LOVPLD using the superPLD^med^ (clone 1-12) variant, which has 4-fold lower activity than superPLD^high^, to investigate if the residual background activity could be eliminated by simply switching to less active superPLD. In line with this hypothesis, LOVPLD(1-12) showed modest activity, about 50% of the original LOVPLD, upon blue light activation **(Fig. 2B)**; however, its background activity was still detectable **(Fig. 2C)**. Excitingly, when LOVPLD was combined with either dimerization system (CRY2–CIBN or iLID), these double-gating constructs exhibited no residual background activity. However, the level of IMPACT labeling was reduced by ∼50% in the iLID system and even further in CRY2–CIBN system, likely due to incomplete dimerization and/or steric hindrance resulting in poor access of active PLD to the membrane. Nevertheless, these examples indicate that LOVPLD can, if desired, be combined with light-induced dimerization systems when further spatiotemporal precision and ultralow background is desired.

Encouraged by these findings, we then evaluated the turn-off kinetics of each of these systems. HEK 293T cells transfected with each construct were exposed to blue light for 30 minutes, and PLD activity was measured by IMPACT labeling after subsequent incubation in the dark for varying durations. Immediately after the illumination period, cells expressing PM-anchored LOVPLD exhibited a 2.2-fold reduction in IMPACT labeling **(Fig. 2D)**. Among the other constructs evaluated, iLID-LOVPLD exhibited an improvement, showing 3.0-fold reduction in activity immediately after the removal of blue light. The difference was even more pronounced after 3 hours, at which point the activity of iLID-LOVPLD has dropped by 23.8-fold, compared to LOVPLD at 4.9-fold. Three hours post-blue light illumination, we found that both iLID-LOVPLD and CRY2-CIBN-LOVPLD returned to baseline levels of IMPACT labeling, whereas LOVPLD and LOVPLD(1-12) still showed substantial residual activity **(Fig. S3A)**. Direct comparison of iLID-LOVPLD and CRY2-CIBN-LOVPLD, the two double-gated systems, indicated faster turn-off kinetics for the former, consistent with the slower dissociation kinetics of the CRY2/CIBN system^35^.

By imaging the localization of iLID-LOVPLD, we observed that it dissociated from the membrane almost immediately in the absence of blue light **(Fig. S3B)**, which we reason contributes to the tighter temporal control of LOVPLD activity in this construct. The success of the double-gated iLID-LOVPLD demonstrates the amenability of LOVPLD to further engineering to produce systems that prioritize high activity or rapid turn-off and highlights its promise as a building block for modular, customizable membrane-editing tools tailored to diverse applications.

### LOVPLD activation in cells specifically impacts PA pools

Having demonstrated precise temporal control of LOVPLD activity using transphosphatidylation-based IMPACT assays, we next evaluated the ability of LOVPLD to produce spatially defined, physiologically relevant PA pools. To visualize the spatial distribution of PA in cells, we used the PA biosensor with superior sensitivity (PASS), a genetically encoded biosensor derived from the Spo20 PA-binding domain^36,37^. We co-expressed GFP-PASS with PM-anchored LOVPLD in HEK 293T cells and used 488 nm laser illumination to visualize PASS localization while activating LOVPLD **(Fig. 3A-D)**. As expected, GFP-PASS exhibited cytosolic localization in cells expressing PM-anchored dLOVPLD regardless of exposure to blue light **(Fig. 3C-D)**. In LOVPLD-expressing cells, however, GFP-PASS recruitment to the PM was observed exclusively after blue light illumination **(Fig. 3A-B)**, despite the constitutive localization of LOVPLD on this membrane, confirming that LOVPLD can selectively produce PA only after its light-induced activation.

**Figure 3.**
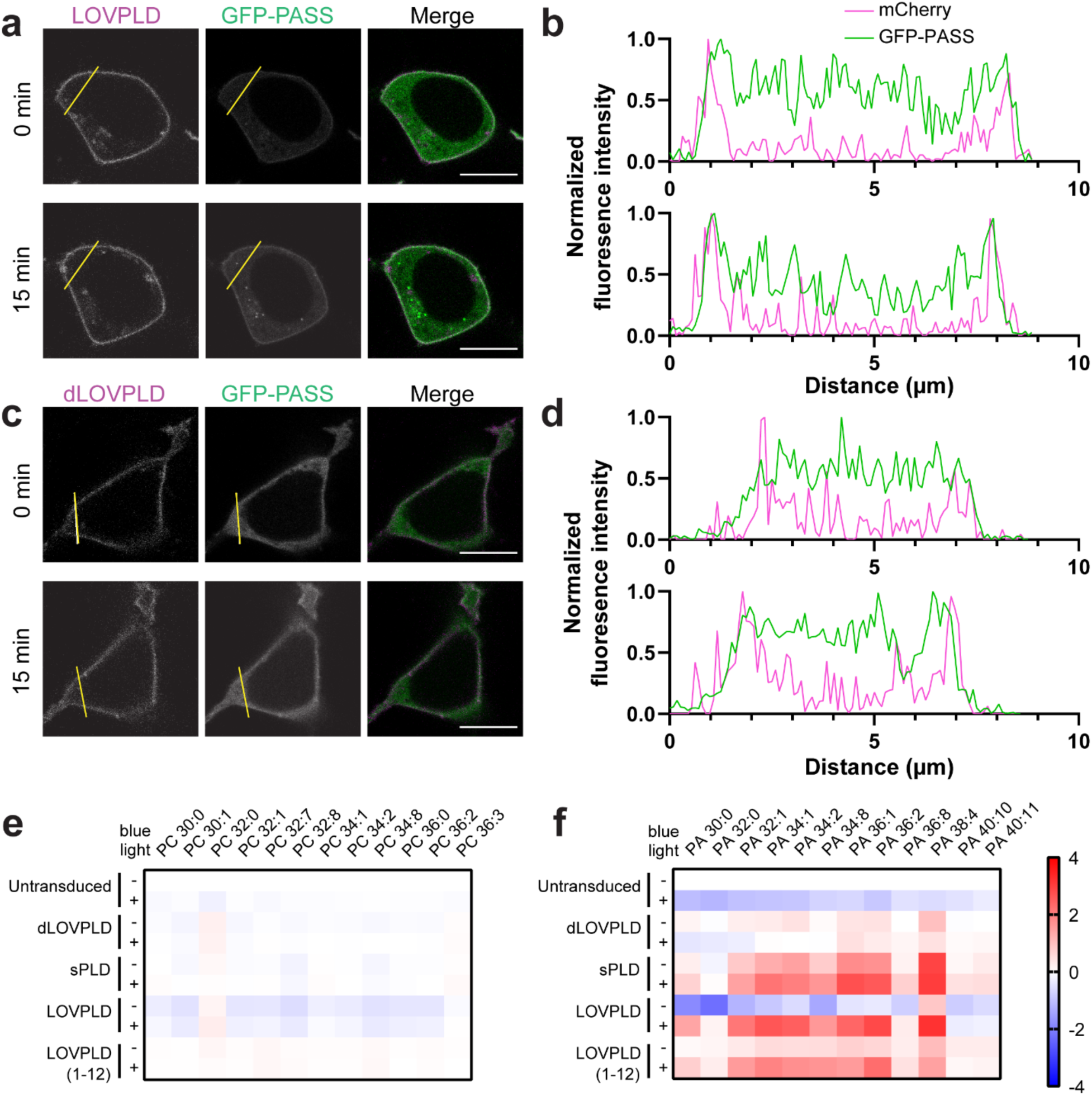
LOVPLD activity has both spatial and substrate specificity. (a–d) Confocal microscopy imaging of the localizations of the PA-binding biosensor GFP-PASS (green) and either PM-anchored LOVPLD (a–b) or dLOVPLD (c–d) (magenta) in HEK 293T cells before (0 min) and after (15 min) blue light illumination, showing that only LOVPLD but not dLOVPLD causes GFP-PASS to deplete from the cytosol and translocate to the PM after blue light illumination. (a and c) Representative single z-slice images. Scale bars: 10 µm. (b and d) Fluorescence intensity profile line plots along yellow lines in (a) and (c). (e–f) Representative results of lipidomics analysis of HEK 293T cells expressing the indicated LOVPLD construct in the absence (–) or presence (+) of blue light stimulation. The relative abundances of major PC (e) and PA (f) species in cellular lipid extracts are displayed as heatmaps. Values are log_2_ fold-change compared to the untransduced –light group (n=3; shown are the 12 most abundant species).

To investigate the identity of the PA species produced by LOVPLD and ascertain whether other major phospholipid populations are affected by LOVPLD activity, we performed lipidomics analyses on cells expressing PM-anchored LOVPLD with or without blue light stimulation. Based on LC–MS analysis of extracted cellular lipids, changes in PC, the major substrate for PLD, were insignificant, as expected due to the high abundance of PC in mammalian cells **(Fig. 3E)**. By contrast, LOVPLD activation resulted in a large increase (10–20 fold) in PA levels, whereas inactive LOVPLD (i.e., kept in the dark) and light-stimulated dLOVPLD did not induce significant changes in PA **(Fig. 3F)**. Cells expressing sPLD also exhibited increased PA levels but as expected, with no light dependency.

An examination of the distribution of individual PA species revealed that activation of LOVPLD at the PM led to increases in several PA species with a trend similar to its parent enzyme superPLD^high^, which mirrors the substrate preference of endogenous PLDs^19,29^, suggesting that the LOV insertion did not alter its acyl chain preference. We also quantified other major classes of phospholipids and found increases in a few PA-derived phospholipid classes upon LOVPLD activation at the PM: LPA, phosphatidylglycerol (PG), and bis(monoacylglycero)phosphate (BMP, also known as lysobisphosphatidic acid) **(Fig. S4A–L)**. Other phospholipid classes, however, including phosphatidylethanolamine (PE), phosphatidylinositol (PI), phosphatidylserine (PS), and lysophospholipids other than LPA, remained largely unchanged. Because LPA and PG are known products of PA metabolism^38–40^, and BMP is a downstream product of PG metabolism (see Discussion), these results indicate that LOVPLD activity is selective for PA production and does not lead to widespread overall changes to the phospholipidome.

Principal component analysis (PCA) also yielded insights into the lipidomics data. The LOVPLD +light conditions clustered with sPLDs, whereas the LOVPLD –light conditions clustered with the catalytically dead controls **(Fig. S4M)**. The results suggest that the features separating samples with and without expected PLD activity were well extracted. Examination of the loading plots revealed that the major contributors to the principal components (PC1 and PC2) that separated the clusters belonged to PA, followed by LPA, PG, and BMP with slightly lower contributions **(Fig. S4N)**. Collectively, these lipidomics analyses establish that LOVPLD can acutely and specifically control production of PA in living cells, with minimal impact on other phospholipid populations.

### LOVPLD reveals organelle-specific differences in PA metabolism

Having thoroughly characterized the properties of PM-targeted LOVPLD, we next examined the feasibility of using LOVPLD to produce PA pools on demand on a variety of intracellular organelle membranes, to address whether acute generation of PA at different subcellular sites would differentially affect phospholipid metabolism. By switching out the membrane anchoring tag, we created LOVPLDs residing on various organelles, including mitochondria, ER, lysosomes, and Golgi, and confirmed their subcellular localization in HEK 293T **(Figs. 4A–D and S5A)** and HeLa cells **(Fig. S5B–E)**. Co-expression of each of these organelle-targeted LOVPLDs with GFP-PASS in HEK 293T cells revealed production of detectable pools of PA on each target membrane within 15 minutes of blue light illumination **(Fig. S6)**. Additionally, IMPACT analysis revealed that LOVPLD targeted to each membrane, with the minor exception of ER-localized LOVPLD, exhibited statistically similar levels of activity **(Fig. 4E)**.

**Figure 4.**
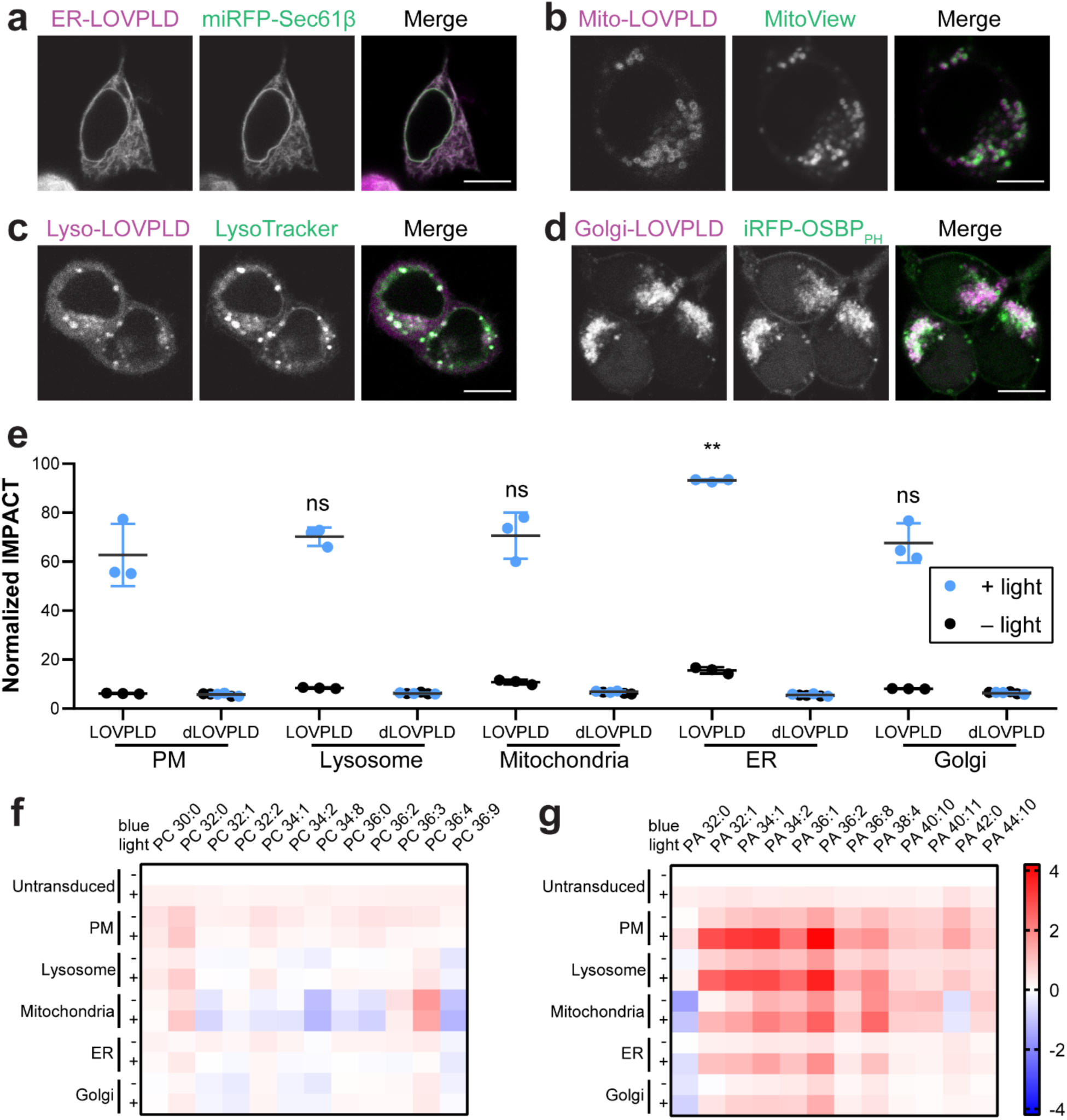
Application of LOVPLDs to probe organelle-selective PA metabolism. (a–d) colocalization of LOVPLD in HEK 293T cells in ER, mitochondria, lysosomes, and Golgi with a) miRFP-Sec61β, b) MitoView 405, c) LysoTracker Red, and d) iRFP-OSBP_PH_. Scale bars: 10 μm. (e) IMPACT labeling and flow cytometry analysis of HEK 293T cells expressing LOVPLD or dLOVPLD targeted to different organelles demonstrates that LOVPLD exhibits similar activity and fold turn-on across different organelles. Shown are mean fluorescence intensities of IMPACT fluorescence in populations of cells expressing the same levels of LOVPLD (assessed by mCherry fluorescence). Statistical significance compared to PM-LOVPLD +light was determined by ordinary one-way analysis of variance (ANOVA) followed by Dunnett’s multiple comparisons test. *, *P* < 0.05; **, *P* < 0.01; ns, not significant. The *P* values for the indicated comparisons are 0.6344, 0.5980, 0.0033, and 0.8697, respectively. (n=3) (f–g) Lipidomics analysis of levels of the 12 most abundant species of PC (f) and PA (g) in HEK 293T cells expressing LOVPLD targeted to different organelle membranes in the presence or absence of blue light illumination. Values are shown as log_2_ fold change compared to the untransduced –light group (n=3).

We reasoned that this consistent expression and activity of the panel of organelle-localized LOVPLDs should enable us to investigate differences in flux through PA metabolism on different organelle membranes. To test this idea, we first analyzed PA levels in cells with LOVPLD activated on different organelles. HEK 293T cells stably expressing LOVPLD at the PM, mitochondria, ER, lysosome, and Golgi complex were generated and exposed to blue light for 30 min, following by analysis of cellular PA levels by LC–MS. Activation of LOVPLD at all organelles had no effect on levels of PC, the abundant phospholipid that is the preferred lipid substrate for PLDs (Fig. 4F). Strikingly, we found that PA abundances were most substantially enriched in cells expressing LOVPLD on the PM and lysosomes, whereas enrichment in mitochondria, the ER, and Golgi compartments was noticeably lower **(Fig. 4G)**.

Because LOVPLD exhibited similar levels of enzymatic activity on these organelles, as revealed by transphosphatidylation-based IMPACT **(Fig. 4E)** and transphosphatidylation tracks well with hydrolysis activity in PLD^PMF^,^20,41^ we can infer that hydrolysis activity, i.e., PA production, should also be similar in all organelles. Therefore, we propose that the lack of PA accumulation on certain organelle membranes is likely due to differential rates of PA consumption on and clearance from these organelles. This hypothesis is further supported by quantification of other phospholipids **(Fig. S7A–L)**. For example, although LOVPLD activation on ER membranes results in lower PA accumulation compared to LOVPLD activation on the PM, increases in PG and BMP abundances are higher for ER samples **(Fig. S7D and G)**.

PCA provided further evidence for differential metabolism of LOVPLD-generated PA on distinct organelles **(Fig. S7M)**. Here, the LOVPLD +light samples clustered distinctly for each organelle and, with the exception of the Golgi complex, were distinct from the control, –light conditions. Similar to the previous analysis of PM-anchored LOVPLD, the loading plots indicated that PA, PG, LPA, and BMP were the major contributors to the principal components (PC1 and PC2) separating the clusters **(Fig. S7N)**.

### LOVPLD reveals effects of organelle-specific PA pools on AMPK signaling

Finally, we set out to harness the unique capabilities of the ultralow background LOVPLDs for manipulating PA-dependent signaling. In contrast to our previous dimerization-based optogenetic PLDs, a distinct advantage of using the ultralow background LOVPLDs for interrogating PA signaling is the tolerance of cells to stable LOVPLD expression and fusion to a wider array of organelle-targeting tags without cytotoxicity from background PA production prior to light stimulus. We envisioned that these advantageous properties would render LOVPLD capable of being applied to assess the contributions of organelle-specific pools of PA toward a sensitive cell signaling pathway.

PA can activate the cellular energy-sensing AMPK signaling pathway through the membrane recruitment of liver kinase B1 (LKB1), which directly phosphorylates and activates AMPK^12^. However, the relationship between PA production at specific membranes and LKB1 localization has not been directly demonstrated, due to the lack of methods to produce PA on a wide array of target membranes. With LOVPLD enabling us to perturb PA production directly and selectively on organelle membranes, we next explored the role of PA in the AMPK signaling pathway. We first tested the ability of LOVPLD-generated PA to induce LKB1 membrane recruitment by co-transfection of HEK 293T cells with LOVPLD and GFP-LKB1 **(Fig. 5A–F)**.

**Figure 5.**
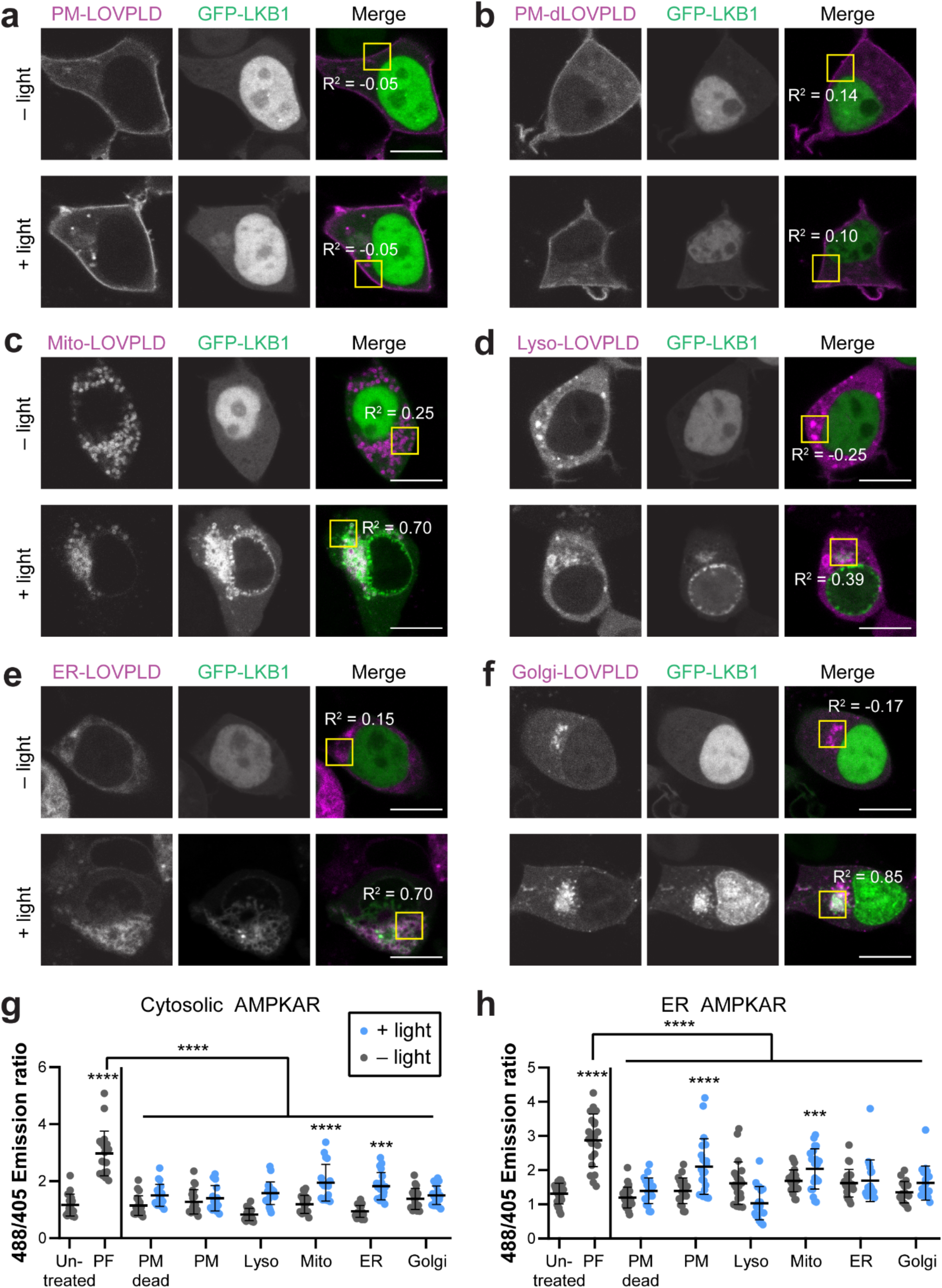
LOVPLD reveals role of PA in recruiting LKB1 and regulating AMPK signaling. (a–f) Acute PA production by organelle-targeted LOVPLDs induces recruitment of GFP-LKB1 to several but not all membranes. The localization of transiently transfected GFP-LKB1 was assessed in HEK 293T cells expressing PM-targeted LOVPLD (a) or dLOVPLD (b), or LOVPLDs targeted to mitochondria (Mito) (c), ER (d), lysosomes (Lyso) (e), or the Golgi complex (f) that were kept in the dark or illuminated with blue light for 30 min. R^2^ values indicate Pearson correlation coefficients of GFP-LKB1 and LOVPLD (mCherry) fluorescence in the yellow boxed region (n=3). Scale bars: 10 µm. (g–h) Imaging of AMPK signaling activity using the ratiometric ExRai-AMPKAR probe expressed either in the cytosol (g) or the cytosolic leaflet of the ER (h) shows increased AMPK activity only when LOVPLDs localized to certain organelles are activated. ExRai-AMPKAR activity was evaluated by assessment of ratio of green emission from 488 nm/405 nm excitation either in the dark (gray) or after 30 min of blue light illumination (blue). PF-06409577 (PF) was used as a positive control for AMPK activation. Asterisks indicate statistical significance compared to untreated cells unless otherwise noted. Statistical significance was determined by ordinary one-way analysis of variance (ANOVA) followed by Tukey’s multiple comparisons test (n=14-24). ****, *P* < 0.0001; ***, *P* < 0.001. The *P* value for ER-LOVPLD + light vs. untreated cells in (g) is 0.0003, and the *P* value for mito-LOVPLD + light vs. untreated cells in (h) is 0.0003.

In control cells, GFP-LKB1 was predominantly nuclear-localized, as expected.^42,43^ Upon blue light illumination, LOVPLD activation on mitochondria, lysosomes, ER, or Golgi membranes induced recruitment of GFP-LKB1 to those organelle membranes, with a concomitant decrease in nuclear LKB1 fluorescence (**Fig. 5C–F**). Importantly, control cells expressing dLOVPLD exhibited no change in GFP-LKB1 localization, and interestingly, we did not observe such GFP-LKB1 recruitment when LOVPLD was expressed on the PM (**Fig. 5A–B**; see Discussion).

We next investigated whether LKB1 recruitment by PA is sufficient to induce local changes in AMPK activity. To assess AMPK activity at the subcellular level, we exploited a recently reported genetically encoded ratiometric fluorescent probe for AMPK activity, ExRai-AMPKAR, which can be targeted to different organelles to report on local AMPK activity following imaging by confocal microscopy.^44^ Using this probe, we investigated whether activation of LOVPLD on different organelle membranes is sufficient to cause local increases in AMPK activity. We first generated ExRai-AMPKAR probes targeted to the cytosol and the cytosolic leaflet of the ER membrane and confirmed their localization by confocal microscopy **(Fig. S8)**. We then co-expressed each ExRai-AMPKAR probe with each of the organelle-targeted LOVPLDs, imaged fluorescence using two excitation wavelengths, 405 nm and 488 nm, and then analyzed changes in 488/405 emission ratios before and after LOVPLD activation. Higher 488/405 emission ratios corresponded to higher levels of AMPK activity, which we validated by using the pharmacological AMPK activator PF-06409577^45^ **(Fig. 5G–H)**.

We observed no change in AMPK activity in cells expressing LOVPLDs without blue light activation, highlighting the importance of the ultralow background LOVPLDs in these studies. Upon blue light activation, cells expressing mitochondrial LOVPLD exhibited increases in AMPK activity in both the cytosol and the ER. Further, cells expressing ER-localized LOVPLD showed a significant increase in cytosolic AMPK activity, and cells expressing PM-localized LOVPLD displayed a significant increase in AMPK activity in the ER. Unexpectedly, these data indicated that increases in AMPK activity did not occur at the organelle membranes where PA was produced. Overall, these results implicate crosstalk between different organelles and interplay between membrane-localized and soluble factors in this signaling pathway, and they underscore the importance of tools with subcellular-level precision for PA production and AMPK visualization for revealing this inter-organelle crosstalk.

## Discussion

In this study, we describe the development and applications of LOVPLD, a photoswitchable membrane editor that catalyzes the conversion of PC to PA and other phospholipids with unprecedented spatiotemporal precision. Compared to our previously reported membrane editors, optoPLDs and optogenetic superPLDs, which use induced proximity to change the localization of enzymes that are constitutively catalytically “on”, LOVPLD features a light-induced turn-on of its catalytic capacity, exhibiting activity in the “on” state that approaches the level of the most active superPLD but a near-baseline background in the “off” state. Manipulation of membrane lipids by LOVPLD offers advantages over conventional techniques such as pharmaceutical activators or inhibitors of lipid biosynthesis pathways and overexpression or knockdown of related enzymes, which almost always involve global manipulation and often manifest on long timescales of hours to days^3,8,18^. Further, the ultralow background activity of LOVPLD opens up previously unattainable applications relative to other optogenetic membrane editors, including studying systems with high sensitivity to PA and those requiring stable expression of the membrane editor.

Our studies using LOVPLD to manipulate cellular PA levels revealed new insights into the metabolic fates of PA. In cells showing high PA accumulation upon organelle targeted LOVPLD activation, we additionally found elevations in levels of LPA and PG, which are products of PA metabolism. Interestingly, we also observed increases in BMP, a low-abundance lipid found mainly on the inner leaflets of late endosomes and lysosomes^46^. Though the full biosynthetic pathway of BMP is still controversial, hydrolysis of PG by a lysosomal phospholipase A (PLA) has been proposed as a first step^47^. As PG abundances increased with LOVPLD activation, our results suggest that the PA–PG–BMP biosynthesis pathway may be one of the earliest cellular responses to overproduction of PA.

Interestingly, even though LOVPLD had similar activity on all organelle membranes tested, we found that LOVPLD activation on ER and Golgi membranes led to much smaller — or undetectable, in the case of the Golgi complex — increases in total cellular PA compared to other organelles (PM, lysosomes, and mitochondria). These results suggest that PA consumption, either via metabolism to other lipids or transport to other organelles, occurs more rapidly on ER and Golgi membranes. The differences in PA consumption between the ER and Golgi complex is particularly noteworthy given the strong coupling between these organelles via vesicular trafficking and non-vesicular pathways. Overall, the lipidomics studies of organelle-selective LOVPLD activation enable us to propose an organelle-based hierarchy for relative rates of PA consumption, with Golgi complex and ER as the sites of most efficient PA metabolism with mitochondria, lysosomes, and PM exhibiting comparatively more sluggish rates of PA metabolism.

Despite the relatively modest accumulation of PA produced by LOVPLDs on some organelle membranes, e.g., ER and mitochondria, such PA pools still elicited biological effects. Notably, PA has been reported to play roles in several major nutrient-sensing pathways, including Hippo and mTOR^10,11,19^. The role of PA in AMPK signaling is comparatively less well defined. The interaction of PA with LKB1 has been suggested to be important, and we had previously observed that cells expressing superPLDs showed slightly elevated global levels of p-AMPK^12,20^. Using an expanded palette of organelle-targeted LOVPLDs whose expression was well tolerated, we first confirmed that PA production on the mitochondria, ER, lysosome, or Golgi complex indeed induces LKB1 recruitment.

Interestingly, PA produced by LOVPLD on the PM did not recruit LKB1, though it was still capable of activating AMPK activity. In epithelial cells, endogenous LKB1 localizes to cell-cell junctions, an interaction that promotes the phosphorylation of LKB1, though the mechanism underlying this localization is unclear^42^. A recent study has proposed an alternate model of PA– AMPK interaction, where PA synthesized on the PM inhibits the biosynthesis of inositol, an AMPK repressor, and thus activates AMPK signaling^48^. Thus, it is possible that though LKB1 is not translocated to the PM via PA recruitment, PA produced on the PM nevertheless still activates AMPK signaling through an LKB1-independent mechanism. Overall, these findings are consistent with previous findings that PA recruits LKB1 and regulates AMPK signaling but also reveal that the underlying mechanisms by which PA activates AMPK signaling may be more complex and/or location dependent^12^.

Our study is also significant for its contribution to the burgeoning field of conditional enzyme activation. Engineering proteins whose functions can allosterically be controlled has emerged as a powerful strategy^49–53^. Our work highlights the potency of direct insertion of an optogenetic conformational switch as a method to directly control enzymatic activity by using light. This approach is fundamentally different from enzymatic activation by translocation, featured in prior optogenetic tools such as optoPLD, as a light-controlled conformational shift achieves rapid and nearly complete enzyme inactivation. Our approach is also different from optogenetic control methods^54,55^ that rely on the steric bulk of the photoactive domain itself to block the enzyme from its substrate to achieve deactivation. Hence, the LOV domain insertion approach — especially when combined with induced proximity modules as in our double-gated iLID-LOVPLD, if necessary — may be more suitable for toxic proteins that cannot be stably expressed, as well as proteins with wide active sites that cannot be easily blocked sterically.

LOV domain insertion is rapidly gaining traction as a general approach for engineering activatable proteins. Most recently, Lee et al. reported LOV-Turbo, a photoswitchable variant of the proximity biotinylation enzyme TurboID that, like LOVPLD, involves interruption of the enzyme sequence with a LOV domain and enables light-mediated enzyme turn-on^53^. LOV domain insertion can also control cellular activity through light-mediated deactivation, rather than activation, of enzymatic activities^51,52^. Further, an alternate optogenetic switch based on the Vivid (VVD) photoreceptor, termed LightR, can activate Src kinases upon light stimulation^56^. Interestingly, the LightR system is relaxed in the dark state and closes when stimulated with blue light, which is opposite of the LOV domain system.

Against this backdrop, our study reporting LOVPLD not only adds to this small but growing collection of photoswitchable functional proteins for probing and manipulating biological systems but also provides unique advances and insights into photoswitchable protein design principles. To our knowledge, LOVPLD is the first demonstration of harnessing photoswitching behavior to control two different enzymatic activities at once, in this case hydrolysis and transphosphatidylation. Exploiting transphosphatidylation as a proxy for hydrolysis activity allowed us to capitalize on the relatively long-lived phosphatidyl alcohols to measure LOVPLD activity when necessary while using the LOVPLDs to interrogate the metabolism and signaling roles of the hydrolysis product, PA. Finally, although we selected a single, optimal insertion site for LOVPLD, our initial screen revealed several additional insertion sites that tolerate LOV domain insertion to some extent, leading to substantial light-dependent activation. We hypothesize that the relatively symmetrical structure of PLD^PMF^ makes it more amenable to such insertion strategies, and it is possible that this property may be translatable to other symmetric enzymes^57^. Overall, this study highlights the utility of LOVPLD as an ultralow background membrane editor, provides new insights into organelle-specific PA metabolism and AMPK signaling, and showcases the potential of optogenetic conformation switches for enabling acute control of metabolic and cell signaling pathways.

## Acknowledgments

This work was supported by an award to J.M.B. from the National Institutes of Health (R01GM151682). X.L. was supported by a Natural Sciences and Engineering Research Council of Canada Postgraduate Scholarship, R.T. was supported by Honjo International, Funai Overseas, and Cornell Fellowships, and M.U. was supported by an Overseas Research Fellowship from the Japan Society for the Promotion of Science and a Long-Term Fellowship from the Human Frontier Science Program. Flow cytometry measurements were performed at the Cornell Institute of Biotechnology Flow Cytometry Facility. We thank the Fromme lab for sharing equipment, Alice Ting for sharing the hLOV1 plasmid, Jin Zhang for sharing the AMPKAR plasmid, and Weizhi Yu for synthesizing the Janelia Fluor 635 HaloTag ligand.

## Author Contributions

R.T. and J.M.B. conceived of the project; X.L., R.T., and J.M.B. designed the study and analyzed data; R.T. and X.L. carried out ConSurf analysis and picked LOV domain insertion sites; M.U. wrote and executed the lipidomics data analysis script and PCA analysis and also assisted in the design of iLID-LOVPLD; X.L. carried out all other experiments; X.L., R.T., M.U., and J.M.B. wrote the manuscript.

