## Supporting Information for "Ultralow background membrane editors for spatiotemporal control of lipid metabolism and signaling"

### Table of Contents

Figure S1. Structural and conservation analysis of sites within PLD chosen for LOV domain insertion ........ S2

Figure S2. Schematic for analysis of LOVPLD activity by IMPACT and generation of HEK 293T cells stably expressing LOVPLD ........ S3

Figure S3. Double-gated optogenetic PLDs exhibit faster turn-off kinetics ........ S4

Figure S4. Lipidomics analysis of PM-targeted LOVPLDs reveals select effects on PA and other phospholipids ........ S5

Figure S5. Targeted LOVPLDs show consistent expression and localization in HEK 293T and HeLa cells ........ S7

Figure S6. Recruitment of the PA biosensor GFP-PASS to organelle membranes by LOVPLD activation ........ S8

Figure S7. Lipidomics analysis of effect of LOVPLDs targeted to multiple organelles ........ S10

Figure S8. Representative images from cytosolic and ER-targeted forms of ExRai-AMPKAR ........ S12

Table S1. Plasmids used for cloning and transfection ........ S13

Table S2. Primers used for cloning ........ S15

Materials and methods ........ S18

References ........ S22

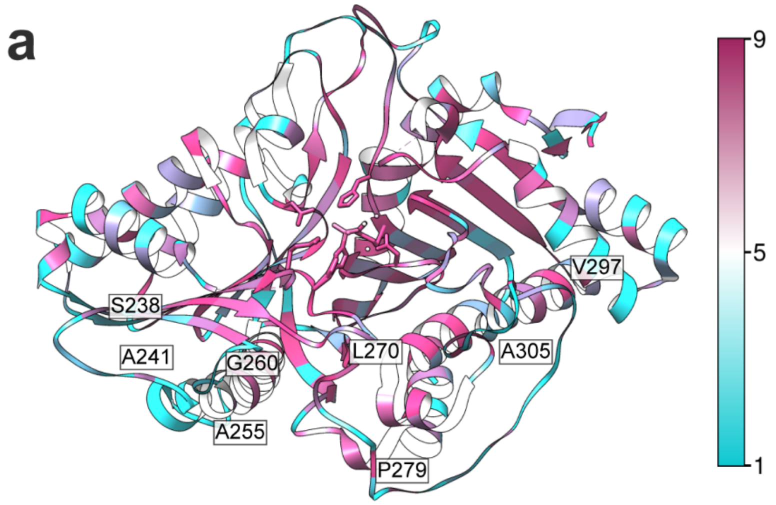


**b**

| LOVPLD variant | AA1 | Conservation score | AA2 | Conservation score |
|---|---|---|---|---|
| A | S38 | 2 | A39 | 1 |
| B | A78 | 9 | T79 | 2 |
| C | G110 | 6 | N111 | 4 |
| D | P123 | 8 | V124 | 1 |
| E | N148 | 1 | I149 | 1 |
| F | G175 | 8 | Q176 | 1 |
| G | A231 | 3 | S232 | 2 |
| H | S238 | 7 | G239 | 1 |
| I | A241 | 4 | G242 | 1 |
| J | A255 | 1 | S256 | 1 |
| K | G260 | 8 | N261 | 1 |
| L | L270 | 9 | G271 | 9 |
| M | P279 | 3 | K280 | 1 |
| N | A290 | 6 | S291 | 1 |
| O | V297 | 1 | G298 | 1 |
| P | A305 | 4 | D306 | 9 |
| Q | K327 | 1 | G328 | 1 |
| R | A361 | 4 | G362 | 6 |
| S | S448 | 3 | S447 | 5 |

**Figure S1. Structural and conservation analysis of sites within PLD chosen for LOV domain insertion.** (a) Evolutionary sequence conservation of $PLD^{PMF}$ was calculated via ConSurf on a scale of 1-9, with 1 being least conserved. Wild type $PLD^{PMF}$ (PDB ID 1V0W, crystal structure shown here) was used as the basis for residue identity and numbering. The eight labeled sites indicate those where LOV domain insertion within $superPLD^{high}$ afforded at least partial blue light activation. (b) Table of LOV domain insertion sites in each of the 19 LOVPLD variants. The LOV domain was inserted in between AA1 and AA2 in $superPLD^{high}$. The conservation score of each AA (amino acid) position is also shown.

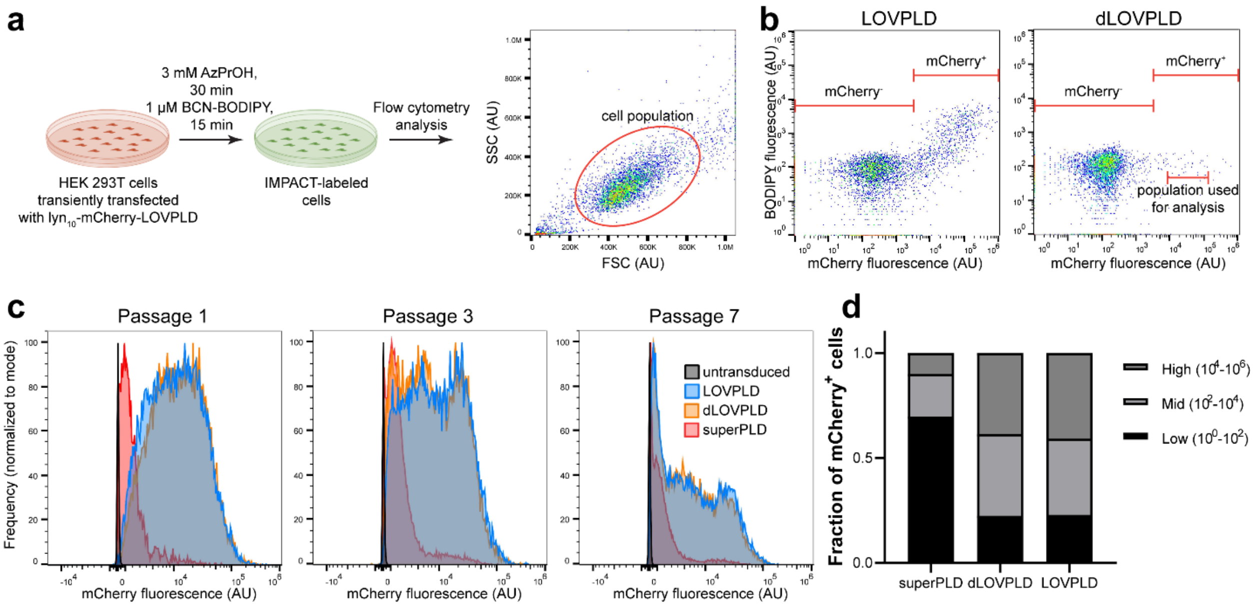


**Figure S2. Schematic for analysis of LOVPLD activity by IMPACT and generation of HEK 293T cells stably expressing LOVPLD.** (a) Schematic showing the workflow of IMPACT labeling and quantification. HEK 293T cells were transiently transfected with the LOVPLD construct. On the day of the experiment, cells were incubated with 3 mM azidopropanol (AzPrOH) for 30 min (with or without blue light illumination). After rinsing with PBS, cells were incubated with 1 µM BCN-BODIPY for 15 min and rinsed again. These labeled cells were then detached from the plate and subjected to flow cytometry analysis, where they were gated based on forward- and side-scatter so that only live, single cells were used for analysis. (b) Representative scatter plots showing IMPACT and LOVPLD (mCherry) fluorescence of cells transfected with LOVPLD or dLOVPLD (catalytically dead LOVPLD) and labeled as in (a). Transfected cells were gated on mCherry expression (mCherry$^+$ corresponding to transfected cells and mCherry$^-$ corresponding to untransfected cells), and the indicated fraction of the mCherry$^+$ population exhibiting moderate levels of mCherry expression was gated ($10^4$–$10^5$ AU). The mean BODIPY fluorescence for this gated group was quantified and used as a measure of IMPACT labeling. (c) HEK 293T cells were transduced with lentiviral constructs encoding either $lyn_{10}$-mCherry-LOVPLD, $lyn_{10}$-mCherry-dLOVPLD, or $lyn_{10}$-mCherry-$superPLD^{high}$. After puromycin selection, cells were passaged every three days, and mCherry levels were quantified via flow cytometry to assess expression of the relevant PLD construct. Note that superPLD was only expressed at extremely low levels, even immediately after transduction and selection (as evidenced by low mCherry fluorescence), whereas LOVPLD and dLOVPLD maintained much higher mCherry fluorescence. (d) Quantification of mCherry$^+$ populations at passage 7 shows that more than 70% of the population for both LOVPLD and dLOVPLD maintained medium to high expression levels, whereas 70% of the cells expressing superPLD do so at very low expression levels.

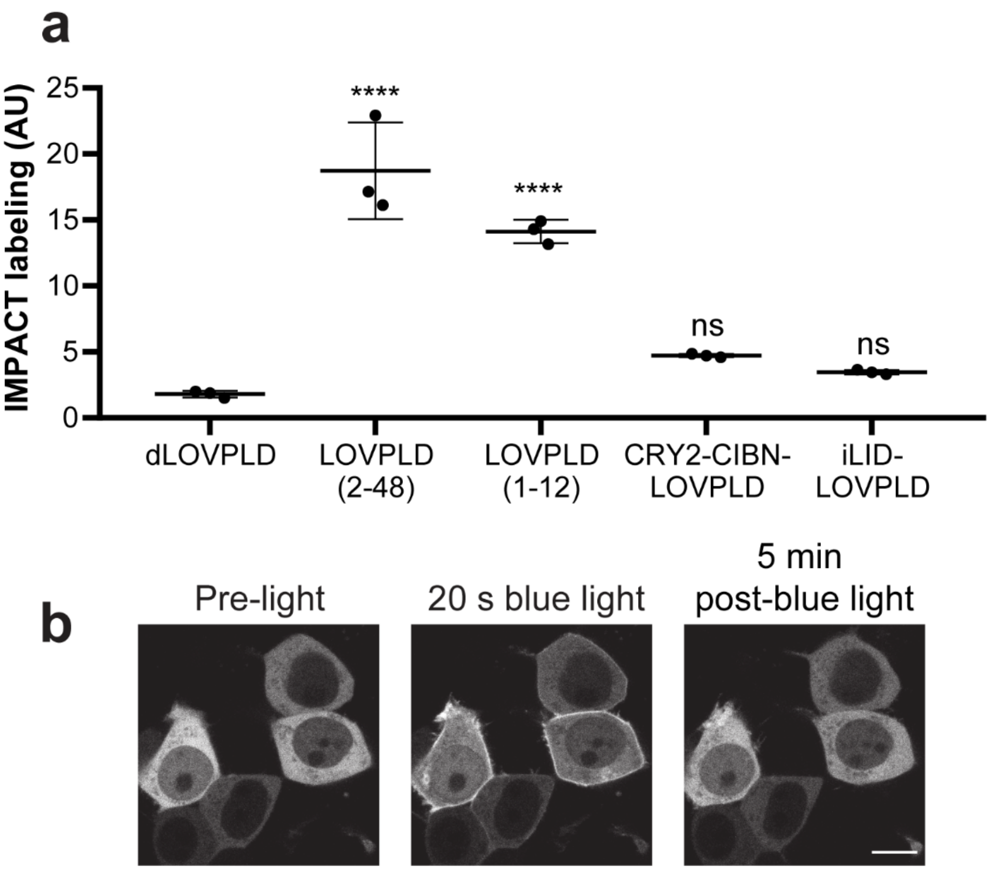


**Figure S3. Double-gated optogenetic PLDs exhibit faster turn-off kinetics.** (a) IMPACT labeling of cells expressing LOVPLD(1-12), CRY2-CIBN-LOVPLD, and iLID-LOVPLD compared to LOVPLD 3 h after removal from blue light stimulation. The activity of the double-gated CRY2-LOVPLD and iLID-LOVPLD both returned to baseline levels, whereas LOVPLD and LOVPLD(1-12) exhibited significant residual activity even after 3 h. Statistical significance compared to dLOVPLD was determined by one-way analysis of variance (ANOVA) followed by Dunnett's multiple comparisons test. ****$P < 0.0001$; NS, not significant. The $P$ values for the indicated comparisons are <0.0001, <0.0001, 0.1729, 0.5881 respectively. (b) Confocal microscopy reveals that iLID-LOVPLD shows cytosolic localization in the dark and is rapidly recruited to the plasma membrane upon blue light illumination. When blue light is removed, iLID-LOVPLD returns to the cytosol. Scale bar: 10 µm.

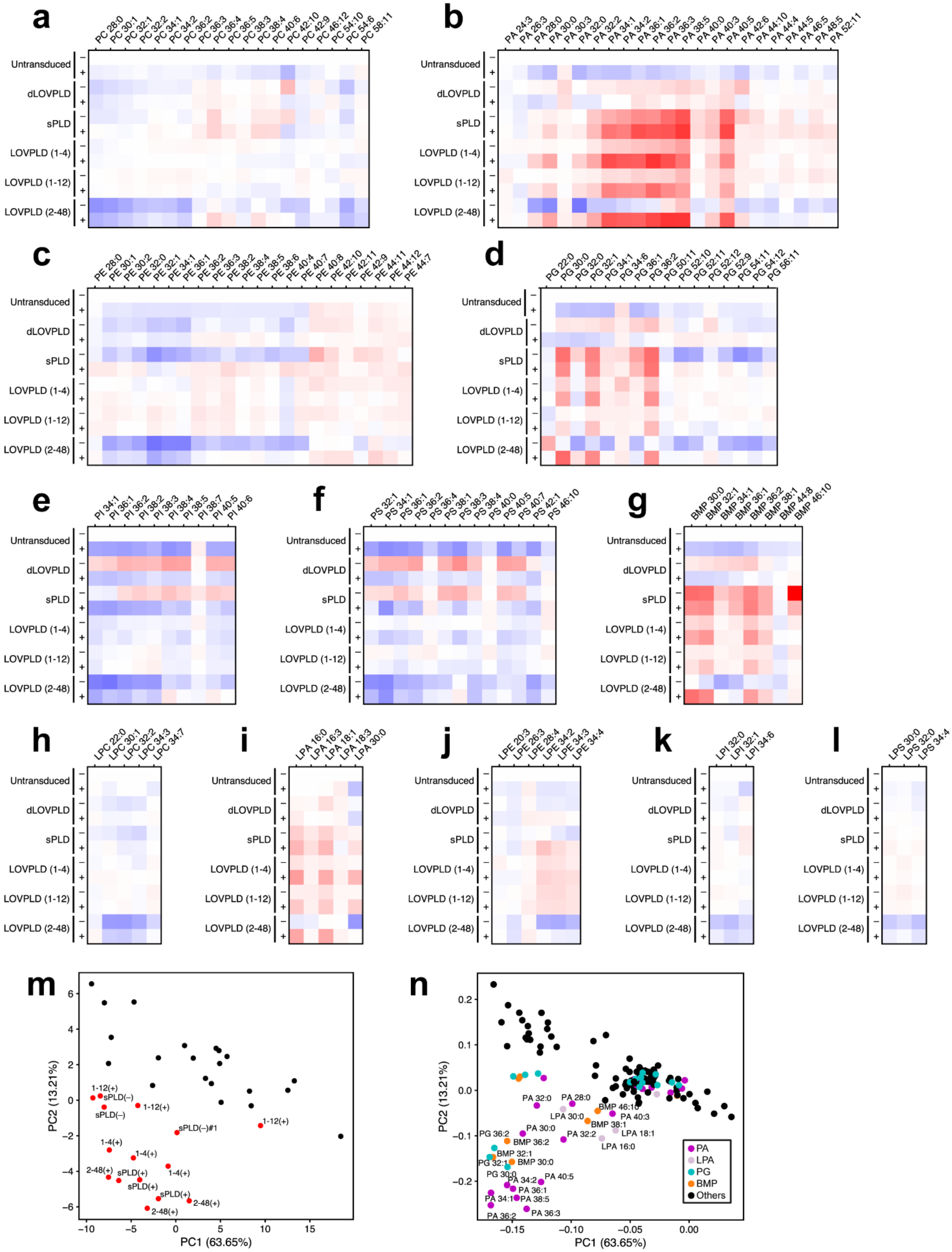
a
PC 28:0 PC 30:1 PC 32:1 PC 32:2 PC 34:1 PC 34:2 PC 36:2 PC 36:3 PC 36:4 PC 36:5 PC 38:3 PC 38:4 PC 40:6 PC 42:10 PC 42:9 PC 46:12 PC 54:10 PC 54:6 PC 58:11
b
PA 24:3 PA 26:3 PA 28:0 PA 30:0 PA 30:3 PA 32:0 PA 32:2 PA 34:1 PA 34:2 PA 36:1 PA 36:2 PA 36:3 PA 38:5 PA 40:0 PA 40:3 PA 40:5 PA 42:6 PA 44:10 PA 44:4 PA 44:5 PA 46:5 PA 48:5 PA 52:11
c
PE 28:0 PE 30:1 PE 30:2 PE 32:0 PE 32:1 PE 34:1 PE 36:1 PE 36:2 PE 36:3 PE 38:2 PE 38:4 PE 38:5 PE 38:6 PE 40:4 PE 40:7 PE 40:8 PE 42:10 PE 42:11 PE 42:9 PE 44:11 PE 44:12 PE 44:7
d
PG 22:0 PG 30:0 PG 32:0 PG 32:1 PG 34:1 PG 34:6 PG 36:1 PG 36:2 PG 50:11 PG 52:10 PG 52:11 PG 52:12 PG 52:9 PG 54:11 PG 54:12 PG 56:11
e
PI 34:1 PI 36:1 PI 36:2 PI 38:2 PI 38:3 PI 38:4 PI 38:5 PI 38:7 PI 40:5 PI 40:6
f
PS 32:1 PS 34:1 PS 36:1 PS 36:2 PS 36:4 PS 38:1 PS 38:3 PS 38:4 PS 40:0 PS 40:5 PS 40:7 PS 42:1 PS 46:10
g
BMP 30:0 BMP 32:1 BMP 34:1 BMP 36:1 BMP 36:2 BMP 38:1 BMP 44:8 BMP 46:10
h
LPC 22:0 LPC 30:1 LPC 32:2 LPC 34:3 LPC 34:7
i
LPA 16:0 LPA 16:3 LPA 18:1 LPA 18:3 LPA 30:0
j
LPE 20:3 LPE 26:3 LPE 28:4 LPE 34:2 LPE 34:3 LPE 34:4
k
LPI 32:0 LPI 32:1 LPI 34:6
l
LPS 30:0 LPS 32:0 LPS 34:4
Untransduced
dLOVPLD
sPLD
LOVPLD (1-4)
LOVPLD (1-12)
LOVPLD (2-48)
–
+
m
PC2 (13.21%)
PC1 (63.65%)
1-12(+)
sPLD(–)
sPLD(–)
1-12(+)
sPLD(–)#1
1-12(+)
1-4(+)
1-4(+)
1-4(+)
2-48(+)
sPLD(+)
sPLD(+)
sPLD(+)
2-48(+)
2-48(+)
n
PC2 (13.21%)
PC1 (63.65%)
PA 32:0
PA 28:0
BMP 46:10
LPA 30:0
PA 40:3
BMP 38:1
PA 30:0
PA 32:2
LPA 18:1
PG 36:2
BMP 36:2
LPA 16:0
BMP 32:1
PG 32:1
BMP 30:0
PG 30:0
PA 34:2
PA 40:5
PA 36:1
PA 34:1
PA 38:5
PA 36:2
PA 36:3
PA
LPA
PG
BMP
Others

**Figure S4. Lipidomics analysis of PM-targeted LOVPLDs reveals select effects on PA and other phospholipids.** Quantification of abundances of all phospholipid classes analyzed in HEK 293T cells expressing plasma membrane-targeted optogenetic PLDs: lyn$_{10}$-mCherry-LOVPLD(2-48), dLOVPLD, constitutively active superPLD$^{high}$ (sPLD), and two LOVPLDs with lower activity backbones (1-4 and 1-12). (a) phosphatidylcholine (PC), (b) phosphatidic acid (PA), (c) phosphatidylethanolamine (PE), (d) phosphatidylglycerol (PG), (e) phosphatidylinositol (PI), (f) phosphatidylserine (PS), (g) bis(monoacylglycero)phosphate (BMP), (h) lysophosphatidylcholine (LPC), (i) lysophosphatidic acid (LPA), (j) phosphatidylethanolamine (LPE), (k) lysophosphatidylinositol (LPI), (l) lysophosphatidylserine (LPS). Abundances are presented as $\log_2$ fold change compared to untransduced HEK 293T cells. (m–n) Principal component analysis of lipidomics data. In the score plot (m), red points indicate conditions where PA production by PLD is expected to occur (i.e., LOVPLDs with light irradiation and sPLD with or without light irradiation) and black points indicate conditions where PA production is not expected to occur (i.e., LOVPLDs without light irradiation and dLOVPLD and untransduced with or without light irradiation. In the loading plot (n), purple, pink, cyan, and orange points indicate PA, LPA, PG, and BMP species, respectively, whereas black points indicate other phospholipid species.

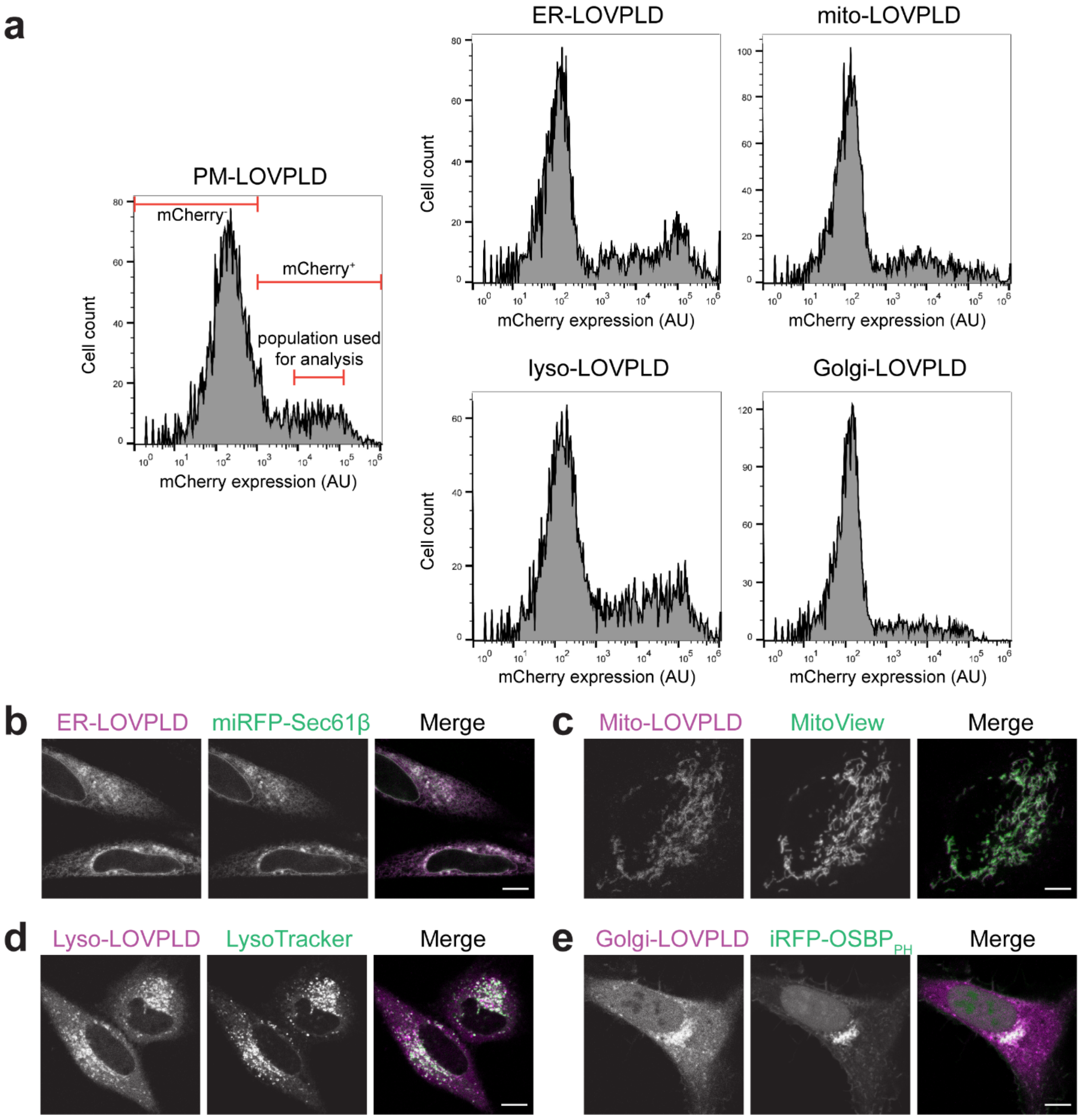


**Figure S5. Organelle-targeted LOVPLDs show consistent expression and localization in HEK 293T and HeLa cells.** (a) Flow cytometry analysis of HEK 293T cells transiently transfected with different organelle-targeted LOVPLDs. LOVPLDs targeted to PM, mitochondria, ER, lysosomes, and the Golgi complex show similar transfection efficiencies and expression levels. The indicated population of mCherry$^+$ cells on the PM-LOVPLD graph ($10^4$–$10^5$ AU) were used to quantify IMPACT labeling. (b–e) Colocalization of organelle markers with LOVPLDs targeted to the ER, mitochondria, lysosomes, and Golgi in HeLa cells. Organelle markers were (b) miRFP-Sec61β, (c) MitoView 405, (d) LysoTracker Red, and (e) iRFP-OSBP$_{PH}$. Scale bars: 10 µm.

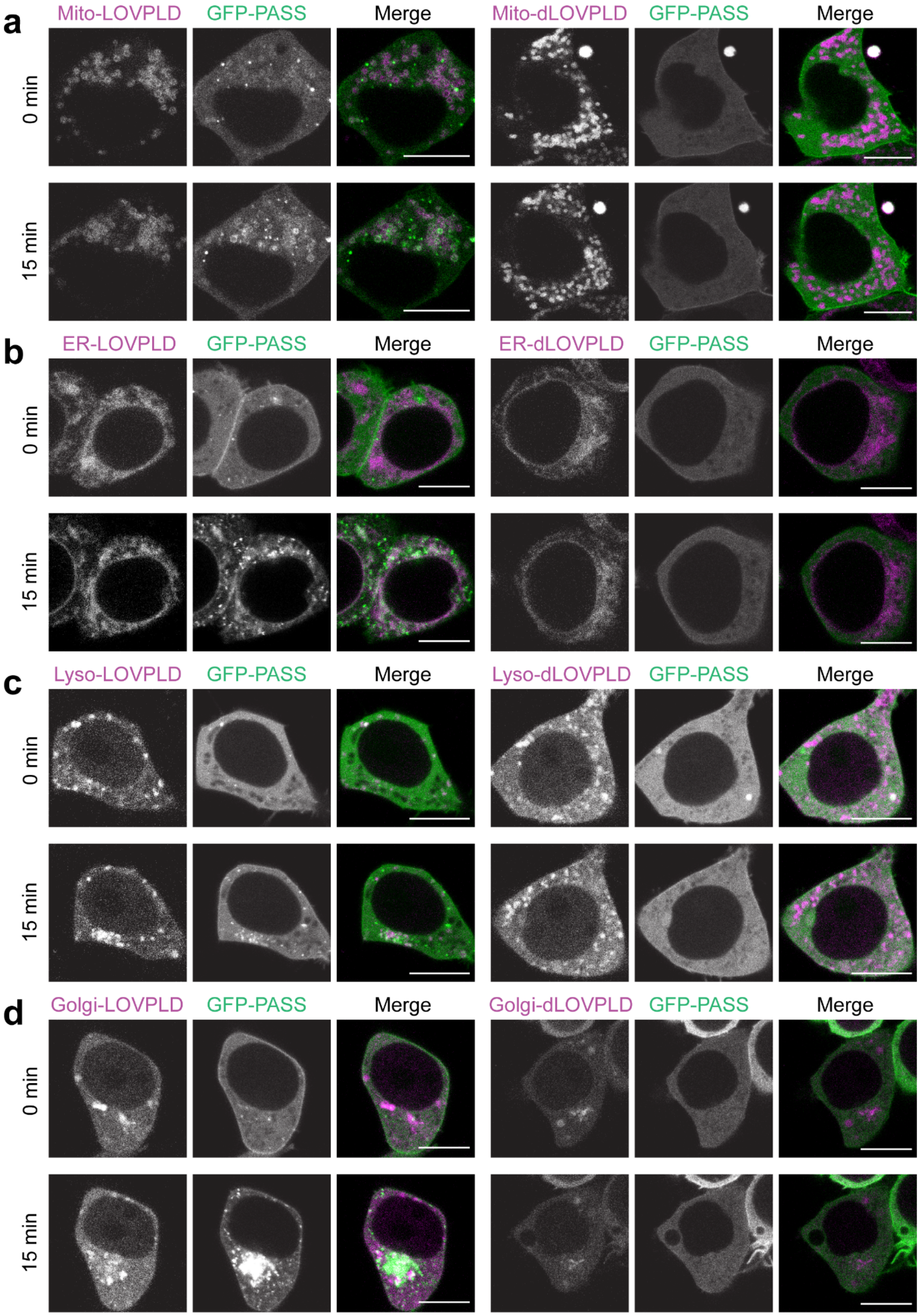
a
Mito-LOVPLD
GFP-PASS
Merge
Mito-dLOVPLD
GFP-PASS
Merge
0 min
15 min
b
ER-LOVPLD
GFP-PASS
Merge
ER-dLOVPLD
GFP-PASS
Merge
0 min
15 min
c
Lyso-LOVPLD
GFP-PASS
Merge
Lyso-dLOVPLD
GFP-PASS
Merge
0 min
15 min
d
Golgi-LOVPLD
GFP-PASS
Merge
Golgi-dLOVPLD
GFP-PASS
Merge
0 min
15 min

**Figure S6. Recruitment of the PA biosensor GFP-PASS to organelle membranes by LOVPLD activation.** HEK 293T cells were co-transfected with GFP-PASS and either LOVPLD or dLOVPLD targeted to (a) mitochondria (Mito), (b) ER, (c) lysosomes (Lyso), and (d) Golgi complex. Cells were imaged by confocal microscopy before (0 min) and after 15 min of intermittent blue light illumination. Scale bars: 10 μm.

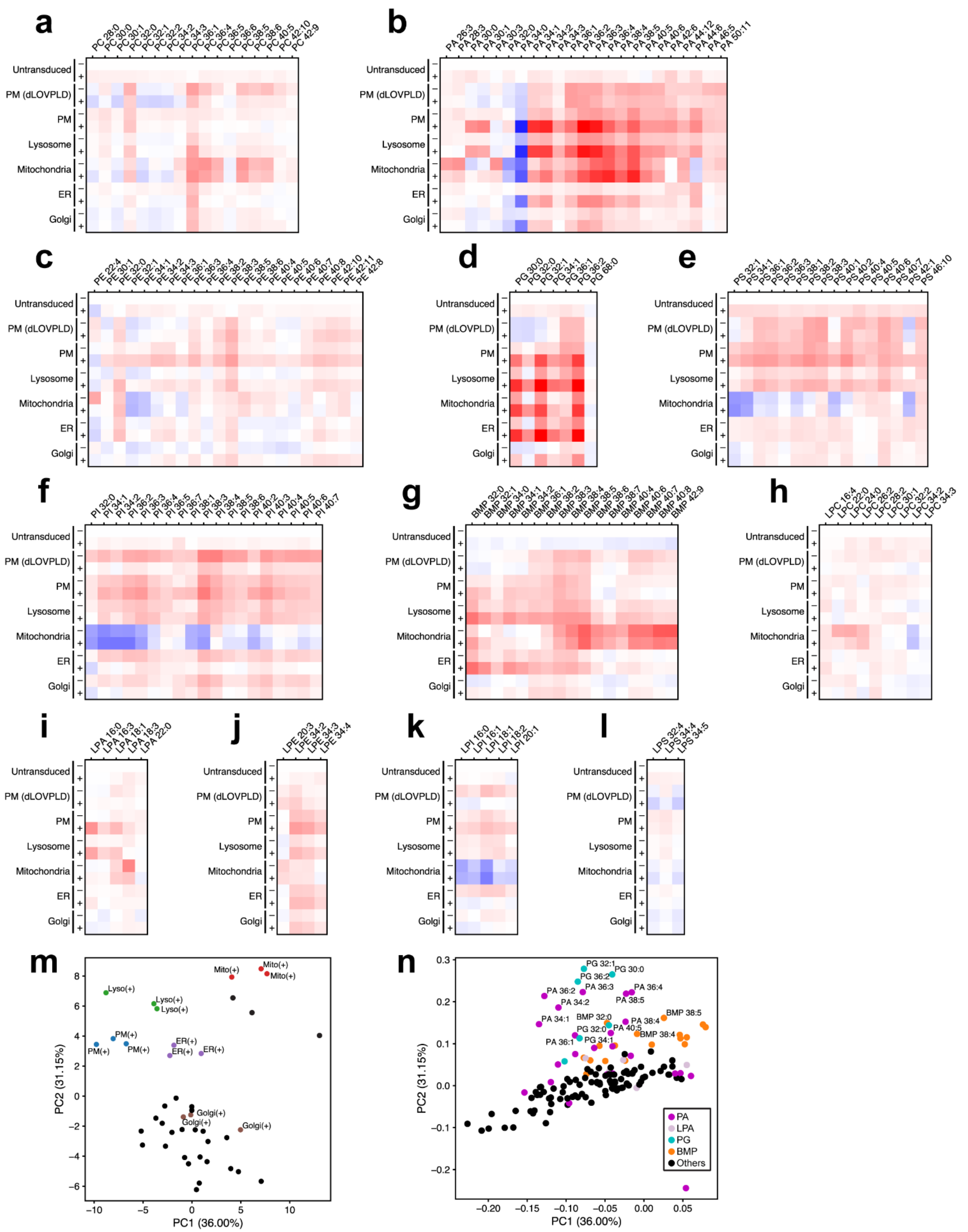
a
PC 28:0 PC 30:0 PC 30:1 PC 32:0 PC 32:1 PC 32:2 PC 34:2 PC 34:3 PC 36:1 PC 36:4 PC 36:5 PC 36:6 PC 38:5 PC 38:6 PC 40:5 PC 42:10 PC 42:9
b
PA 26:3 PA 28:3 PA 30:0 PA 30:1 PA 30:3 PA 32:0 PA 34:0 PA 34:1 PA 34:2 PA 34:3 PA 36:1 PA 36:2 PA 36:3 PA 36:4 PA 38:4 PA 38:5 PA 40:5 PA 40:6 PA 42:6 PA 44:12 PA 44:6 PA 46:5 PA 50:11
c
PE 22:4 PE 30:1 PE 32:0 PE 32:1 PE 34:1 PE 34:2 PE 34:3 PE 36:1 PE 36:3 PE 36:4 PE 38:2 PE 38:3 PE 38:5 PE 38:6 PE 40:4 PE 40:5 PE 40:6 PE 40:7 PE 40:8 PE 42:10 PE 42:11 PE 42:8
d
PG 30:0 PG 32:0 PG 32:1 PG 34:1 PG 36:1 PG 36:2 PG 68:0
e
PS 32:1 PS 34:1 PS 36:1 PS 36:2 PS 36:3 PS 38:1 PS 38:2 PS 38:3 PS 40:1 PS 40:2 PS 40:4 PS 40:5 PS 40:6 PS 40:7 PS 42:1 PS 46:10
f
PI 32:0 PI 34:1 PI 34:2 PI 36:2 PI 36:3 PI 36:4 PI 36:5 PI 36:7 PI 38:1 PI 38:3 PI 38:4 PI 38:5 PI 38:6 PI 40:2 PI 40:3 PI 40:4 PI 40:5 PI 40:6 PI 40:7
g
BMP 32:0 BMP 32:1 BMP 34:0 BMP 34:1 BMP 34:2 BMP 36:1 BMP 38:2 BMP 38:3 BMP 38:4 BMP 38:5 BMP 38:6 BMP 38:7 BMP 40:4 BMP 40:6 BMP 40:7 BMP 40:8 BMP 42:9
h
LPC 16:4 LPC 22:0 LPC 24:0 LPC 26:2 LPC 28:2 LPC 30:1 LPC 32:2 LPC 34:2 LPC 34:3
i
LPA 16:0 LPA 16:3 LPA 18:1 LPA 18:3 LPA 22:0
j
LPE 20:3 LPE 34:2 LPE 34:3 LPE 34:4
k
LPI 16:0 LPI 16:1 LPI 18:1 LPI 18:2 LPI 20:1
l
LPS 32:4 LPS 34:4 LPS 34:5
Untransduced
PM (dLOVPLD)
PM
Lysosome
Mitochondria
ER
Golgi
−
+
m
Mito(+) Mito(+) Mito(+)
Lyso(+) Lyso(+) Lyso(+)
PM(+) PM(+) PM(+)
ER(+) ER(+) ER(+)
Golgi(+) Golgi(+) Golgi(+)
PC1 (36.00%)
PC2 (31.15%)
n
PG 32:1 PG 30:0 PG 36:2 PA 36:2 PA 36:3 PA 36:4 PA 38:5 PA 34:2 BMP 38:5 BMP 32:0 PA 38:4 PA 34:1 PG 32:0 PA 40:5 BMP 38:4 PA 36:1 PG 34:1
PA
LPA
PG
BMP
Others
PC1 (36.00%)
PC2 (31.15%)

**Figure S7. Lipidomics analysis of effect of LOVPLDs targeted to multiple organelles.** Quantification of abundances of all phospholipid classes analyzed in HEK 293T cells expressing LOVPLDs targeted to the PM, lysosomes, mitochondria, ER, and Golgi. (a) phosphatidylcholine (PC), (b) phosphatidic acid (PA), (c) phosphatidylethanolamine (PE), (d) phosphatidylglycerol (PG), (e) phosphatidylinositol (PI), (f) phosphatidylserine (PS), (g) bis(monoacylglycero)phosphate (BMP), (h) lysophosphatidylcholine (LPC), (i) lysophosphatidic acid (LPA), (j) phosphatidylethanolamine (LPE), (k) lysophosphatidylinositol (LPI), (l) lysophosphatidylserine (LPS). Abundances are presented as $\log_2$ fold change compared to untransduced HEK 293T cells. (m–n) Principal component analysis of lipidomics data. In the score plot (m), organelle LOVPLDs +light samples are labeled and shown in color. In the loading plot (n), purple, pink, cyan, and orange points indicate PA, LPA, PG, and BMP species, respectively, and black points indicate other phospholipid species.

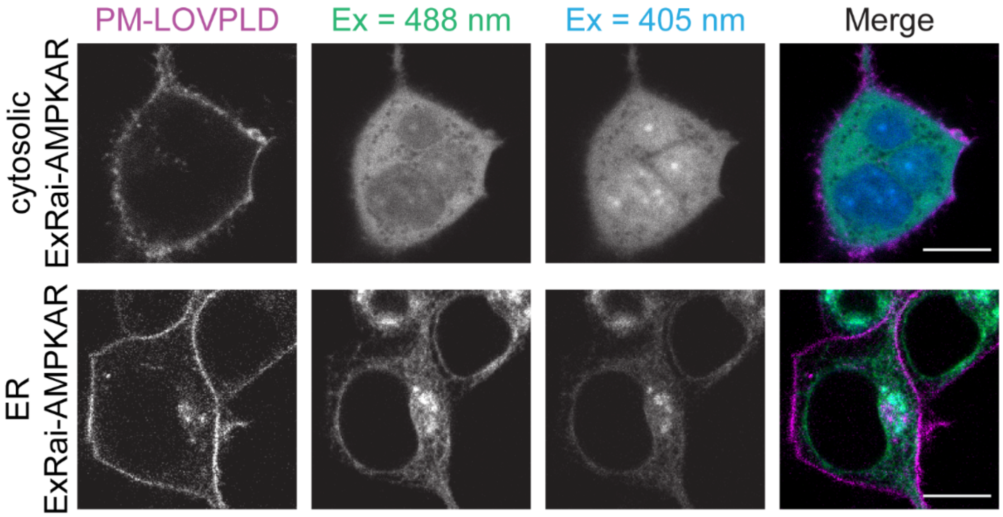


**Figure S8. Representative images from cytosolic and ER-targeted forms of ExRai-AMPKAR.** Confocal microscopy images of ExRai-AMPKAR expressed either in the cytosol or the ER, showing fluorescence emission in the 500-600 nm range when excited with either 488 nm or 405 nm light. The ratio of fluorescence emission from excitation at both wavelengths (488/405) was calculated for each cell and used to quantify AMPK activity as shown in **Fig. 5**. Scale bars: 10 μm.

**Table S1.** Plasmids used for cloning and transfection.

| | Plasmid | Source | Notes |
|---|---|---|---|
| 1 | CRY2-mCherry-superPLD$^{high}$-P2A-CIBN-CAAX | Tei et al. 2023[1] | used as template for cloning |
| 2 | CRY2-mCherry-superPLD$^{med}$-P2A-CIBN-CAAX | Tei et al. 2023[1] | used as template for cloning |
| 3 | CRY2-mCherry-superPLD$^{low}$-P2A-CIBN-CAAX | Tei et al. 2023[1] | used as template for cloning; Fig 1d |
| 4 | AKAP1-hLOV-TEVcs-GAL4bd-VP64-V5 | Coukos et al. 2021[2] | used as template for cloning |
| 5 | Lyn$_{11}$-GFP-CAAX | Idevall-Hagren et al. 2012[3] | used as template for cloning |
| 6 | pCAGGS-PIF3-mEGFP | Uda et al. 2017[4] | used as template for cloning |
| 7 | pLL7.0-tgRFPt-SSPB WT | Guntas et al. 2015[5] | used as template for cloning |
| 8 | iRFP-OSBP1$_{PH}$ | P. De Camilli (Yale University, New Haven, CT) | used as template for cloning; Fig 4a |
| 9 | C1(1-29)-TurboID-V5-pCDNA3 | Branon et al. 2018[6] | used as template for cloning |
| 10 | pLX304 CMV HA-HaloTag-FRB-sTurboID (C) | Cho et al. 2020[7] | used as template for cloning |
| 11 | pCDNA3-lyn$_{10}$-mCherry-LOVPLD-A | This study | Fig 1c |
| 12 | pCDNA3-lyn$_{10}$-mCherry- LOVPLD-B | This study | Fig 1c |
| 13 | pCDNA3-lyn$_{10}$-mCherry- LOVPLD-C | This study | Fig 1c |
| 14 | pCDNA3-lyn$_{10}$-mCherry- LOVPLD-D | This study | Fig 1c |
| 15 | pCDNA3-lyn$_{10}$-mCherry- LOVPLD-E | This study | Fig 1c |
| 16 | pCDNA3-lyn$_{10}$-mCherry- LOVPLD-F | This study | Fig 1c |
| 17 | pCDNA3-lyn$_{10}$-mCherry- LOVPLD-G | This study | Fig 1c |
| 18 | pCDNA3-lyn$_{10}$-mCherry- LOVPLD-H | This study | Fig 1c |
| 19 | pCDNA3-lyn$_{10}$-mCherry- LOVPLD-I | This study | Fig 1c |
| 20 | pCDNA3-lyn$_{10}$-mCherry- LOVPLD-J | This study | Fig 1c |
| 21 | pCDNA3-lyn$_{10}$-mCherry- LOVPLD-K | This study | Fig 1c |
| 22 | pCDNA3-lyn$_{10}$-mCherry- LOVPLD-L | This study | Fig 1c |
| 23 | pCDNA3-lyn$_{10}$-mCherry- LOVPLD-M | This study | Fig 1c |
| 24 | pCDNA3-lyn$_{10}$-mCherry- LOVPLD-N | This study | Fig 1c |
| 25 | pCDNA3-lyn$_{10}$-mCherry- LOVPLD-O | This study | Fig 1c |
| 26 | pCDNA3-lyn$_{10}$-mCherry- LOVPLD-P (LOVPLD) | This study | Fig 1c-d, 2b-d, 3a, 5a, 5g-h, S2a, S3a, S6, S8 |
| 27 | pCDNA3-lyn$_{10}$-mCherry- LOVPLD-Q | This study | Fig 1c |
| 28 | pCDNA3-lyn$_{10}$-mCherry- LOVPLD-R | This study | Fig 1c |
| 29 | pCDNA3-lyn$_{10}$-mCherry- LOVPLD-S | This study | Fig 1c |
| 30 | pCDNA3-lyn$_{10}$-mCherry-superPLD | This study | Fig 1c |
| 31 | pcDNA3-lyn$_{10}$-mCherry-dsPLD | This study | Fig 1c |
| 32 | pCDNA3-lyn$_{10}$-mCherry-dLOVPLD | This study | Fig 1c-d, 2b-d, 3a, 5a, 5g-h, S2a, S3a, S6, S8 |

| | | | |
|---|---|---|---|
| 33 | PJL143-pcDNA3-CRY2-mCherry-LOVPLD-P2A-CIBN-CAAX | This study | Fig 2b-d, S3a |
| 34 | PJL144-pCAGGS-lyn$_{10}$-hLOV-ssrA-P2A-LOVPLD-mCherry-sspB | This study | Fig 2b-d, S3a-b |
| 35 | pCDNA3-lyn$_{10}$-mCherry-LOVPLD (1-12 backbone) | This study | Fig 2b-d, S3a |
| 36 | EGFP-PASS | Zhang et al. 2014[8] | Fig 3a, S7 |
| 37 | pCDH1-lyn$_{10}$-mCherry-dLOVPLD | This study | Fig 3b-e, S4, S5 |
| 38 | pCDH1-lyn$_{10}$-mCherry-superPLD | This study | Fig 3b-e, S4, S5 |
| 39 | pCDH1-lyn$_{10}$-mCherry-LOVPLD | This study | Fig 3b-e, S4, S5 |
| 40 | pCDH1-lyn$_{10}$-mCherry-LOVPLD (1-12 backbone) | This study | Fig 3b-e, S4, S5 |
| 41 | pCDH1-lyn$_{10}$-mCherry-LOVPLD (1-4 backbone) | This study | Fig S4, S5 |
| 42 | iRFP-Sec61β | P. DeCamilli (Yale University, New Haven, CT) | Fig 4a |
| 43 | pCDNA3-AKAP1-mCherry-LOVPLD | This study | Fig 4a, 4e, 5c, 5g-h, S6 |
| 44 | pCDNA3-AKAP1-mCherry-dLOVPLD | This study | Fig 4e, 5c, 5g-h, S6 |
| 45 | pCDNA3-C1-mCherry-LOVPLD | This study | Fig 4b, 4e, 5d, 5g-h, S6 |
| 46 | pCDNA3-C1-mCherry-dLOVPLD | This study | Fig 4e, 5d, 5g-h, S6 |
| 47 | pCDNA3-p18-mCherry-LOVPLD | This study | Fig 4c, 4e, 5e, 5g-h, S6 |
| 48 | pCDNA3-p18-mCherry-dLOVPLD | This study | Fig 4e, 5e, 5g-h, S6 |
| 49 | pCDNA3-LOVPLD-mCherry-OSBP$_{PH}$ | This study | Fig 4d, 4e, 5f, 5g-h, S6 |
| 50 | pCDNA3-dLOVPLD-mCherry-OSBPPH | This study | Fig 4e, 5f, 5g-h, S6 |
| 51 | pCDNA3-p18-HaloTag-LOVPLD | This study | Fig 4c |
| 52 | pCDH1-AKAP1-mCherry-LOVPLD | This study | Fig 4f-g |
| 53 | pCDH1-AKAP1-mCherry-dLOVPLD | This study | Fig 4f-g |
| 54 | pCDH1-C1-mCherry-LOVPLD | This study | Fig 4f-g |
| 55 | pCDH1-C1-mCherry-dLOVPLD | This study | Fig 4f-g |
| 56 | pCDH1-p18-mCherry-LOVPLD | This study | Fig 4f-g |
| 57 | pCDH1-p18-mCherry-dLOVPLD | This study | Fig 4f-g |
| 58 | pCDH1-LOVPLD-mCherry-OSBP$_{PH}$ | This study | Fig 4f-g |
| 59 | pCDH1-dLOVPLD-mCherry-OSBP$_{PH}$ | This study | Fig 4f-g |
| 60 | GFP-LKB1 | Tei et al. 2023[1] | Fig 5a-g |
| 61 | pCDNA3-ExRai-AMPKAR | Schmitt et al. 2022[9] | Fig 5g, S8 |
| 62 | pCDNA3-Tex264tm-ExRai-AMPKAR | This study | Fig 5h, S8 |

**Table S2.** Primers used for cloning.

| | **Name** | **Sequence** |
|---|---|---|
| 1 | HindIII-Kozak-mCherry-S | GTCAAGCTTGCCACCATGGTGAGCAAG |
| 2 | BamHI-mCherry-A | GTCGGATCCCTTGTACAGCTCGTCCATGC |
| 3 | EcoRI-hLOV-S | GTCAGAATTCGAGTTTCGGGCAACCACAC |
| 4 | NotI-hLOV-A | GTCAGCGGCCGCAGCAATCTGAAAGGCTGTCTTTTTG |
| 5 | pcDNA3-lyn10-S | GAGACCCAAGCTTGCCACCATGGGATGTATT |
| 6 | $lyn_{10}$-KpnI-A | GTTATCCTCCTCGCCCTTGCTCACGGTACCCGAACCTGAGCTACCG |
| 7 | BamHI-PLD-S | GTCGGATCCGCAGATTCAGCTACACCACATTTG |
| 8 | EcoRI-PLDS41-A | GTCAGAATTCACTTCCGTCCAATTTATTTCCAC |
| 9 | EcoRI-PLDA81-A | GTCAGAATTCGGCATTCCCTATATTTTCAGTCATC |
| 10 | EcoRI-PLDG113-A | GTCAGAATTCGCCTCTAGCTGCGGAC |
| 11 | EcoRI-PLDP126-A | GTCAGAATTCGGGTGCAGCCCCAAC |
| 12 | EcoRI-PLDN151-A | GTCAGAATTCGTTTTCTGCGGCTTTGC |
| 13 | EcoRI-PLDG178-A | GTCAGAATTCACCAGTAAGGGCGCTTTG |
| 14 | EcoRI-PLDA234-A | GTCAGAATTCAGCGATGTTACTCTTGTTTTGGC |
| 15 | EcoRI-PLDS241-A | GTCAGAATTCGGAAGCTGCAAACCAGAC |
| 16 | EcoRI-PLDA244-A | GTCAGAATTCTGCGTTGCCGGAAGC |
| 17 | EcoRI-PLDA258-A | GTCAGAATTCTGCCTTGGGGTTTGTATCTTTATG |
| 18 | EcoRI-PLDG263-A | GTCAGAATTCCCCGGTGGCAGGTGATG |
| 19 | EcoRI-PLDL273-A | GTCAGAATTCCAACCCTCCAACGGCTATTATTG |
| 20 | EcoRI-PLDP282-A | GTCAGAATTCGGGGTCCACATCTTTGATGC |
| 21 | EcoRI-PLDA296-A | GTCAGAATTCAGCTGTTGGCAGATCGG |
| 22 | EcoRI-PLDV304-A | GTCAGAATTCTACCACACACTTAGTATCGCTAG |
| 23 | EcoRI-PLDA312-A | GTCAGAATTCGGCGTTAGTATTGTCGTGAAG |
| 24 | EcoRI-PLDK327-A | GTCAGAATTCTTTTGCTGAAGCAACAAGAGCTC |
| 25 | EcoRI-PLDA368-A | GTCAGAATTCAGCAGCCATTTTAGCAGCC |
| 26 | EcoRI-PLDS456-A | GTCAGAATTCTGAGTCGACGGAAACCAACTTATG |
| 27 | ApaI-TTA-PLD-A | GTCAGGGCCCTTAAGCGTTGCAGATTCCTCTTG |
| 28 | NotI-A42PLD-S | GTCAGCGGCCGCTGCCGCTGATCCCAGTG |
| 29 | NotI-T82PLD-S | GTCAGCGGCCGCTACCAGAACCGTAGATATTAGTACAC |
| 30 | NotI-N114PLD-S | GTCAGCGGCCGCTAATAAGTTGAAAGTTAGGATACTGG |
| 31 | NotI-V127PLD-S | GTCAGCGGCCGCTGTCTACCACATGAATGTAATACCTTC |
| 32 | NotI-I152PLD-S | GTCAGCGGCCGCTATAACTTTGAACGTGGCTAGTATGAC |
| 33 | NotI-G179PLD-S | GTCAGCGGCCGCTGGAATAAATAGCTGGAAGGATGATTAC |
| 34 | NotI-S235PLD-S | GTCAGCGGCCGCTTCCGTCTGGTTTGCAGC |
| 35 | NotI-G242PLD-S | GTCAGCGGCCGCTGGCAACGCAGGGTGC |
| 36 | NotI-gly245PLD-S | GTCAGCGGCCGCTGGGTGCATGGCCACTATG |
| 37 | NotI-S259PLD-S | GTCAGCGGCCGCTTCACCTGCCACCGGG |
| 38 | NotI-N264PLD-S | GTCAGCGGCCGCTAATGTCCCAATAATAGCCGTTG |

| | | |
|---|---|---|
| 39 | NotI-G274PLD-S | GTCAGCGGCCGCTGGCGTTGGCATCAAAGATG |
| 40 | NotI-K283PLD-S | GTCAGCGGCCGCTAAGTCCACTTTCAGGCCC |
| 41 | NotI-S297PLD-S | GTCAGCGGCCGCTAGCGATACTAAGTGTGTGGTAG |
| 42 | NotI-G305PLD-S | GTCAGCGGCCGCTGGACTTCACGACAATACTAACGC |
| 43 | NotI-D313PLD-S | GTCAGCGGCCGCTGATAGAGACTATGATACAGTCAACCC |
| 44 | NotI-S328PLD-S | GTCAGCGGCCGCTAGTCACATTGAAATATCTCAGCAG |
| 45 | NotI-G369PLD-S | GTCAGCGGCCGCTGGGGTTAAGGTGAGAATTGTGG |
| 46 | NotI-S457PLD-S | GTCAGCGGCCGCTTCCACATTTTATATCGGTTCAAAAAAT TTG |
| 47 | PLDH167A-S | AAAAACAGCTTTCTCCTGGAACGCTAGCAAAATCTTGGT TGTAGAC |
| 48 | PLDH167A-A | GTCTACAACCAAGATTTTGCTAGCGTTCCAGGAGAA AGCTGTTTTT |
| 49 | HindIII-Kozak-AKAP1-S | GTCAAGCTTGCCACCATGATGGCCATCCAGCTGC |
| 50 | AgeI-AKAP1-A | GTCACCGGTGGTGGCCACGGGAG |
| 51 | HindIII-Kozak-p18-S | GTCAAGCTTGCCACCATGATGGGGTGCTGCTACAG |
| 52 | AgeI-p18-A | GTCACCGGTGTAGTTGGGCTCGGCTC |
| 53 | HindIII-Kozak-C1-S | GTCAAGCTTGCCACCATGATGGACCCTGTGGTCGT |
| 54 | AgeI-C1-A | GTCACCGGTTGATCCCCCGCCTCC |
| 55 | pcDNA3-HindIII-Kozak-BamHI-PLDN-S | ACTCACTATAGGGAGACCCAAGCTTGCCACCATGGGATC CGCAGATTCAGCTAC |
| 56 | mCherry-KpnI-PLDN-A | TCGCCCTTGCTCACGGTACCAGCGTTGCAGATTCCTCTT G |
| 57 | KpnI-linker-mCh-S | GTCGGTACCGGAGGCGGTTCAGGCG |
| 58 | AgeI-linker-mCh-A | GTCACCGGTTCCTCCTGCTCCCGCTG |
| 59 | AgeI-OSBP1$_{PH}$-S | GTCACCGGTGGCTCGGCTCGAGAGG |
| 60 | ApaI-OSBP1$_{PH}$-A | GTCAGGGCCCTCATGCCAGCATCTTCACAG |
| 61 | KpnI-HaloTag-S | GTCGGTACCATGGCAGAAATCGGTACTGG |
| 62 | BsrGI-HaloTag-A | GTCTGTACAGGCCGGAAATCTCGAGCG |
| 63 | pCDH1-XbaI-HindIII-Kozak-lyn$_{10}$-S | TTTGACCTCCATAGAAGATTCTAGAAGCTTGCCACCATG GGATG |
| 64 | pCDH1-AsiSI-ApaI-TTA-PLD-A | CTGACGGGCACCGGAGCGATCGCGGGCCCTTAAGCGTT GCAG |
| 65 | XbaI-Kozak-p18-S | GTCATCTAGAGCCACCATGATGGGGTGC |
| 66 | XbaI-Kozak-AKAP1-S | GTCATCTAGAGCCACCATGATGGCCATCC |
| 67 | XbaI-Kozak-C1-S | GTCATCTAGAGCCACCATGATGGCCATC |
| 68 | XbaI-Kozak-BamHI-PLD-S | GTCATCTAGAGCCACCATGGGATCCGC |
| 69 | linker-BamHI-PLD-S | GGAGCAGGTCGATCGGGATCCGCAGATTCAGCTACACC AC |

| | | |
|---|---|---|
| 70 | linker-EcoRI-PLD-A | GTAGCGCCGCTTCCGAATTCAGCGTTGCAGATTCCTCTT G |
| 71 | NotI-mCherry-S | TCCGGAGGTTCCGGCGGCCGCATGGTGAGCAAGGGCGA G |
| 72 | XbaI-mCherry-A | CCCGCTGCTGCGCCTCTAGACAGCTCGTCCAT |
| 73 | EcoRI-KS-PLD-S | ATCATTTTGGCAAAGAATTCGCCACCATGG |
| 74 | NheI-linker-A | GCTAGCACCAGCACTATCTCTAGAT |
| 75 | NheI-hLOV-S | AGTGCTGGTGCTAGCTTTCGGGC |
| 76 | AflII-SsrA-A | GCCGCTTCCCTTAAGAAAGTAATTTTCGTCGTTC |
| 77 | XbaI-hLOV-S | GCTGTCTAGAGGTGGATCTGGAGGTTCAGGTGGAAGTG CTAGCTTTCGGGCAACCACAC |
| 78 | SalI-SsrA-hLOV-A | ATCCGTCGACTTAAAAGTAATTTTCGTCGTTCGCTGCCT CAGCAATCTGAAAGGCTG |
| 79 | PLD-NheI-S | CCGATCTGCCAACAGCCAGCGATA |
| 80 | linker-XhoI-PLD-A | CGGAACCTCCGGATCCTCCCTCGAGAGCGTTGCAGATT CC |
| 81 | AflII-P2A-S | CTTAAGGGAAGCGGCGCTACA |
| 82 | PLD-NheI-A | ACACACTTAGTATCGCTGGCTGTTG |
| 83 | EGFP-XbaI-linker-S | GGCATGGACGAGCTGTCTAGAGGCGCAGCAGCGGGAG |
| 84 | EcoRV-linker-A | CATGATATCAGATCCGCCCGATCGA |
| 85 | EcoRV-SspB-S | GGATCTGATATCATGAGCTCCCCGAAACGCC |
| 86 | SalI-SspB-A | AGGGAAAAAGATCCGTCGACTAACCAATATTCAGCTCG |
| 87 | AgeI-KpnI-PLD-S | ATTGACCGGTACCATGGCAGATTCAGCT |
| 88 | L-XhoI-PLD-A | CGGAACCTCCGGATCCTCCCTCGAGAGCGTTGCAGATT CC |
| 89 | AgeI-ExRai-S | GTCACCGGTATGAGGAGAGTGGCTACCC |
| 90 | ApaI-TTA-ExRai-A | GTCAGGGCCCTTAGCGATCAACTTTGTTCTGC |
| 91 | HindIII-TEX264$_{TM}$-S | GTCAAGCTTGCCACCATGTCGGACCTGCTAC |
| 92 | AgeI-TEX264$_{TM}$-A | GTCACCGGTCCCAGCCAGTAGCCCTG |

### Materials and Methods

#### Plasmids and cloning

All primers used for cloning can be found in Supplementary Table 1. To generate mCherry-LOVPLD, mCherry was first amplified by PCR from CRY2-mCherry-superPLD$^{high}$-P2A-CIBN-CAAX using primers 1-2 and digested with HindIII and BamHI restriction enzymes, and then ligated into a HindIII/BamHI-digested pCDNA3 vector backbone to create pCDNA3-mCherry. hLOV1 was amplified by PCR from AKAP1-hLOV-TEVcs-GAL4bd-VP64-V5 (Addgene #170995) using primers 3-4 and inserted into pCDNA3-mCherry using EcoRI/NotI restriction sites. Lyn$_{10}$ was amplified by PCR from Lyn$_{11}$-GFP-CAAX (obtained from Pietro De Camilli, Yale University) using primers 5-6 and inserted into pCDNA3-mCherry-hLOV using megaprimer PCR.

To clone the LOVPLD candidates A-S, primers 8-26 were used in conjunction with primer 7 to amplify 19 PLD fragments from pCDNA-CRY2-mCherry-superPLD$^{high}$-P2A-CIBN-CAAX, which were digested with BamHI and EcoRI restriction sites and inserted into BamHI/EcoRI-digested pCDNA3-Lyn$_{10}$-mCherry-hLOV backbone. Primers 28-46 were then used in conjunction with primer 27 to amplify 19 fragments corresponding to the C terminal half of PLD, which were digested with NotI and ApaI restriction enzymes and inserted into a plasmid backbone created from the previous step containing the corresponding N-terminal PLD fragment. PM-anchored non-optogenetic superPLD$^{high}$ (sPLD) was cloned by PCR amplification of superPLD$^{high}$ using primers 7 and 27 and insertion into pCDNA3-Lyn$_{10}$-mCherry-hLOV backbone using BamHI and ApaI restriction sites. dsPLD and dLOVPLD were cloned by site-directed mutagenesis using primers 47-48 on sPLD and LOVPLD P respectively.

Mitochondria, lysosome, and ER-targeting versions of LOVPLD were cloned by swapping out Lyn$_{10}$ for AKAP1 (amplified by PCR from AKAP1-hLOV-TEVcs-GAL4bd-VP64-V5 using primers 49-50), p18 (ATGGGGTGCTGCTACAGCAGCGAGAACGAGGACTCGGACCAGGACCGAGAGGAGC GGAAGCTGCTGCTGGACCCTAGCAGCCCCCCTACCAAAGCTCTCAATGGAGCCGAGC CCAACTAC (sequence provided by Pietro De Camilli, Yale University), amplified by PCR using primers 51-52), and C1 (amplified by PCR from C1(1-29)-TurboID-V5-pCDNA3 (Addgene #107173)) using primers 53-54) respectively using HindIII/AgeI restriction sites. To clone Golgi-targeting LOVPLD, mCherry (amplified by PCR from CRY2-mCherry-superPLD$^{high}$-P2A-CIBN-CAAX using primers 55-56) and LOVPLD (amplified by PCR using primers 57-58) were first ligated using overlap PCR, and the PH domain of OSBP1 was amplified by PCR from iRFP-OSBP1$_{PH}$ (a gift from Pietro De Camilli, Yale University) using primers 59-60. The overlap PCR product was digested with HindIII and AgeI, and OSBP$_{PH}$ was digested with AgeI and ApaI restriction enzymes, and the two fragments were inserted into a HindIII/ApaI-digested pCDNA3 backbone to create LOVPLD-mCherry-OSBP$_{PH}$. Lysosome-targeting LOVPLD with HaloTag was cloned by swapping out mCherry for HaloTag (PCR amplified from pLX304 CMV HA-HaloTag-FRB-sTurboID (Addgene #153003) using primers 61-62) using KpnI and BsrGI restriction sites.

PM-targeted LOVPLD for lentivirus production was cloned by PCR amplification of pCDNA3-Lyn$_{10}$-mCherry-LOVPLD using primers 63-64 and insertion into XbaI/AsiSI-digested pCDH1 vector using Gibson assembly. To make lentiviral versions of the other targeted LOVPLDs, mitochondria, lysosome, and ER-targeted LOVPLDs were amplified by PCR using primers 65-66 in conjunction with primer 64, and Golgi-targeted LOVPLD amplified using primers 68 and 60, and fragments were then inserted into an XbaI/ApaI digested pCDH1-Lyn$_{10}$-mCherry-LOVPLD backbone. CRY2-CIBN-LOVPLD was cloned by PCR amplification of

LOVPLD using primers 69-70 and insertion into BamHI/EcoRI-digested CRY2-mCherry-PLD-P2A-CIBN-CAAX backbone using Gibson assembly. To clone iLID-LOVPLD, mCherry was amplified by PCR using primers 71-72, $Lyn_{10}$ using primers 73-74, and iLID(hLOV-ssrA) using primers 75-78 all from pCDNA3-$Lyn_{10}$-mCherry-LOVPLD; the P2A peptide was amplified by PCR using primers 79-80, PLD using primers 81-82, and a linker using primers 83-84 all from pCDNA-CRY2-mCherry-superPLD$^{high}$-P2A-CIBN-CAAX; and sspB was PCR amplified from pLL7.0-tgRFPt-SSPB (Addgene #60415) using primers 85-86. All fragments were inserted into an AflII/SalI-digested pCAGGS backbone (pCAGGS-PIF3-mEGFP, Addgene #100283) using Gibson assembly, and LOVPLD was amplified using primers 87-88 and inserted using KpnI and XhoI restriction sites.

ER-targeted ExRai-AMPKAR is cloned by PCR amplification of ExRai-AMPKAR from pCDNA3-ExRai-AMPKAR (Addgene #192446) using primers 89-90 and the transmembrane domain of TEX264 (ATGTCGGACCTGCTACTACTGGGCCTGATTGGGGGCCTGACTCTCTTACTGCTGCTGACGCTGCTGGCCTTTGCCGGGTACTCAGGGCTACTGGCTGGG, sequence predicted based on cDNA of HEK 293T cells using a hidden Markov model[10]) using primers 91-92 inserting into a pCDNA3 backbone digested using HindIII/AgeI/ApaI.

**Mammalian cell culture, transfection, and lentiviral transduction**

Cells were grown in DMEM (Corning) supplemented with 10% FBS (Corning) and 1% penicillin/streptomycin (Corning) at 37 °C in a 5% $CO_2$ atmosphere. For poly-L-lysine pre-treatment, plates were treated with 0.1 mg/mL poly-L-lysine (Sigma Aldrich; P2636) in PBS for 20 min at 37 °C, followed by triple rinses with PBS.

For transient transfection, HEK 293T cells or HeLa cells were transfected using PEI MAX (Polysciences; 24765-100). Briefly, plasmids were pre-mixed with PEI MAX in growth media (1 µg plasmid and 5 µg PEI MAX per 2 mL DMEM) and added to cells 16-24 h after seeding. The cells were then covered with aluminum foil to keep cells in the dark and incubated for another 16–24 h before being labeled or analyzed.

For lentivirus production, HEK 293TN cells seeded on a 6-well plate were incubated in Transfectagro (Corning) supplemented with 10% FBS containing plasmids pre-mixed with Lipofectamine 2000 (Thermo Fisher 11668019) using 0.5 µg envelope plasmid, 1.5 µg packaging plasmid, 1.5 µg LOVPLD plasmid, and 6 µL Lipofectamine 2000 per 35-mm well of a 6-well plate. 16 h after transfection, the transfection media was replaced with regular DMEM media, and media was collected every 8 h after transfection to obtain virus-containing media. For lentiviral transduction, HEK 293T cells seeded on a 6-well plate (pre-treated with poly-L-lysine) were incubated in 1.5 mL virus-containing media supplemented with 0.5 mL fresh media and 0.8 µg/mL polybrene (Millipore Sigma). The 6-well plate was covered with aluminum foil. After 24 h, virus-containing media was replaced with fresh DMEM media, and cells were incubated in the dark for another 24 h before analysis.

**Blue light illumination setup**

A homemade light box was built by attaching six strips of dimmable, 12 V blue-LED tape light (1000Bulbs.com; 2835–60-IP65-B1203) on the inside of a plastic tray. For optogenetics experiments, the light box was placed inside the $CO_2$ incubator using an AC Outlet Power Bank (Omars; 24,000 mAh, 80 W) as a power supply. An outlet timer (BN-LINK) was used to switch the light on and off automatically to enable 3-s intervals of blue light in every 1 min.

**IMPACT labeling and flow cytometry analysis**

HEK 293T cells transfected with LOVPLD constructs were treated with 3 mM azidopropanol (AzPrOH) for 30 min at 37 °C under intermittent blue light illumination (3-s pulses every 1 min). For figure 1D, 500 μM AzPrOH was used instead. After three rinses with PBS, cells were treated with 1 μM bicyclononyne-BODIPY fluorophore[11] (BCN-BODIPY) for 15 min at 37 °C. Cells were again rinsed three times with PBS then trypsinized, fixed in 4% paraformaldehyde (Electron Microscopy Sciences), and analyzed using a Attune NxT cytometer (Thermo Fisher). Acquired data were analyzed using the Python package FlowCytometryTools and FlowJo.

**Confocal microscopy**

HEK 293T cells or HeLa cells were seeded on 35-mm glass-bottom imaging dishes (MatTek Corporation or Matsunami) pre-coated with poly-L-lysine and transfected as described above. For IMPACT labeling, cells were treated with 3 mM AzPrOH in growth media for 30 min at 37°C with or without light illumination as above and rinsed three times with PBS. Then the cells were treated with 1 μM BCN-BODIPY for 15 min at 37°C, rinsed another three times with PBS, and then incubated in growth media for 15 min before imaging. For GFP-PASS and GFP-LKB1 colocalization, the dish was placed in an imaging chamber kept at 37°C and 5% $CO_2$ atmosphere, and images were taken every 30 s using 488 and 561 laser illumination. For ExRai-AMPKAR quantification, dishes were incubated in an external incubator for 30 min with or without blue light and then transferred to the imaging chamber and imaged. Images were acquired using Zeiss Zen Blue 2.3 on a Zeiss LSM 800 confocal laser scanning microscope equipped with Plan Apochromat objectives (40× 1.4 NA) and two GaAsP detectors. Solid-state lasers (405, 488, 561,and 640 nm) were used to excite fluorophores. Acquired images were processed using ImageJ.

**Live cell staining**

HEK 293T cells were seeded on 35-mm glass-bottom imaging dishes pre-coated with poly-L-lysine and transfected with organelle-targeted LOVPLDs as described above. For mitochondria-targeted LOVPLD, cells were transfected with AKAP1-mCherry-LOVPLD 20 h prior to imaging and incubated with MitoView 405 (Biotium) in growth media for 15 min; cells were imaged after replacing the media with fresh DMEM. For lysosome-targeted LOVPLD, cells were transfected with p18-HaloTag-LOVPLD 20 h prior to imaging. Prior to imaging, cells were incubated with 1 μM Janelia Fluor 635 HaloTag ligand[12] for 10 min, then replaced with 50 nM LysoTracker Red DND-99 (Invitrogen) and incubated for 10 min; cells were imaged after replacing the media with fresh DMEM.

**LC–MS**

HEK 293T cells were seeded in 12-well plates and transduced with lentivirus as described above. 36-48 h after transduction, 1 μM puromycin was added to the growth media to select for transduced cells. After 24 h, cells were incubated for 30 min at 37 °C with or without light illumination and then rinsed once with PBS and subjected to Bligh-Dyer lipid extraction. Briefly, 250 μL methanol, 125 μL acetic acid (20 mM in water), and 100 μL PBS were added to each well, and cells were scraped and transferred into 1.5 mL Eppendorf tubes. 500 μL chloroform was then added to each tube, and the solution was mixed thoroughly by shaking vigorously for 5 min. The tubes were then centrifuged briefly to facilitate phase separation. The bottom organic layer was collected, transferred into new tubes, and dried under a steady stream of nitrogen gas. For injection into the LC–MS, the dried lipid film were resuspended in 150 μL chloroform/methanol/water mix (73:23:3), filtered through a 0.45 μm syringe filter (Millipore Sigma), and 1-3 μL was injected for analysis. LC–MS analysis was performed on an Agilent 6230 electrospray ionization–time-of-flight MS coupled to an Agilent 1260 HPLC equipped with a Luna 3 μm Silica LC Column (Phenomenex; 50 × 2 mm) using a binary gradient elution system where solvent A was chloroform/methanol/ammonium hydroxide (85:15:0.5) and solvent B was chloroform/methanol/water/ammonium hydroxide (60:34:5:0.5). Separation was achieved using a linear gradient from 100% A to 100% B over 10 min. Phospholipid species were detected using an Agilent Jet Stream source operating in negative mode. LC–MS data was acquired using Agilent Mass Hunter Workstation software (Version B.08.00).

**Lipidomics data analysis**

LC–MS data was exported as .mzdata.xml format using Mass Hunter Qualitative Analysis Navigator (Version B.08.00). Non-targeted lipidomics data analysis was performed with custom Python scripts. First, a target file containing m/z values was generated based on the formula of each lipid species, and retention times were manually set for each class of lipids. Based on this target file, mass spectra of each lipid were extracted according to the assigned retention time. Targets whose mass spectral peaks were not within ±10 ppm of the theoretical values in the untransduced, –light condition were excluded from subsequent analysis. Finally, retention times to be integrated were manually adjusted for each lipid species to obtain areas under the curve (AUCs). For each lipid species, obtained AUCs were first normalized by calculating $\log_2$(fold change) against the average values of the untransduced, –light conditions. Data were then normalized by subtracting the average values for each lipid species throughout the samples, followed by PCA analysis using scikit-learn version 1.1.1 with Python 3.8.13.

**Statistical methods**

For experiments involving quantification of comparisons between more than two independent groups, statistical significance was determined by ordinary one-way analysis of variance (ANOVA) followed by Tukey's multiple comparisons test or Dunnett's multiple comparisons test, performed in GraphPad Prism (version 8.0.1 for Windows). Colocalization was calculated in FIJI using the Coloc 2 plugin to determine Pearson correlation coefficients. All experiments were performed in at least three biological replicates on different days. Exact numbers of replicate experiments and sample sizes are provided in each figure legend.

**Safety statement**

No unexpected or unusually high safety hazards were encountered.
